# Oxytocin signals via Gi and Gq to drive persistent CA2 pyramidal cell firing and strengthen CA3-CA1 neurotransmission

**DOI:** 10.1101/2020.05.07.082727

**Authors:** Katherine W. Eyring, Jingjing Liu, Gabriele M. König, Shizu Hidema, Katsuhiko Nishimori, Evi Kostenis, Richard W. Tsien

## Abstract

The oxytocin receptor (OXTR) is concentrated in specific brain regions, exemplified by hippocampal subregion CA2, that support social information processing. Oxytocinergic modulation of CA2 directly affects social behavior, yet how oxytocin regulates activity in CA2 remains incompletely understood. We found that OXTR stimulation acts via closure of M-current potassium channels in all OXT-sensitive CA2 neurons. M-current inhibition was persistent in CA2 pyramidal cells, whose prolonged burst firing required functional coupling of the OXTR to both Gαq and Gαi proteins. Other neuromodulators acted via distinct patterns of G-protein signaling to induce CA2 pyramidal neuron burst firing, underscoring its likely importance. CA2 burst firing impacted hippocampal subregion CA1 where *stratum oriens*-resident CA1 interneurons were targeted more strongly than CA1 pyramidal cells. Oxytocinergic modulation of interneurons, via CA2 pyramidal cell input and directly, triggered a long-lasting enhancement of CA3-CA1 transmission. Thus, transient activation of oxytocinergic inputs may initiate long-lasting recording of social information.

## Introduction

Understanding how neuromodulators shape neural activity and relate to neuropsychiatric disease are cornerstones of modern neuroscience. Oxytocin, a peptide hormone with well-defined roles in parturition and lactation, has been identified as a modulator of prosocial behavior across the animal kingdom (Donaldson & Young, 2008; Garrison et al., 2012; Gimpl & Fahrenholz, 2001). Increased levels of oxytocin are associated with trust, generosity and facial recognition in humans (Bartz, Zaki, Bolger, & Ochsner, 2011; Kosfeld, Heinrichs, Zak, Fischbacher, & Fehr, 2005; Skuse et al., 2014; Zak, Stanton, & Ahmadi, 2007), while variations in its receptor and plasma levels have been observed in patients with autism spectrum disorder (ASD)(LoParo & Waldman, 2014; Modahl et al., 1998; Wu et al., 2005). Oxytocin treatment of ASD and other disorders with atypical social behavior has shown variable but encouraging preclinical promise in patients and animal models (Feifel et al., 2010; Penagarikano et al., 2015; Young & Barrett, 2015), prompting further investigation into how oxytocin signals in the brain. A more refined mechanistic understanding of central oxytocin signaling might facilitate the development of more precise and effective therapies.

Expression of the oxytocin receptor (OXTR) is concentrated in specific regions of the brain (Insel & Shapiro, 1992), highlighting potential hubs of social information processing. Coupled with use of transgenic mouse lines (Hidema et al., 2016; Nakajima, Gorlich, & Heintz, 2014; Yoshida et al., 2009), the development of the first specific OXTR antibody (Mitre et al., 2016) has enabled experiments to define what cell-types respond to oxytocin and model how neural circuits underlying social behavior are modulated (Menon et al., 2018; Oettl et al., 2016; Tirko et al., 2018; Xiao, Priest, Nasenbeny, Lu, & Kozorovitskiy, 2017). The hippocampal sub-region CA2, which is distinguished from neighboring areas CA1 and CA3 by its anatomy, physiology and gene expression profile (Dudek, Alexander, & Farris, 2016), is enriched with OXTRs (Lee, Caldwell, Macbeth, Tolu, & Young, 2008; Mitre et al., 2016; Tirko et al., 2018). There, pyramidal cells express the OXTR (Tirko et al., 2018) and their activity is required for the encoding, consolidation and recall of a short-term social memory in mice (Hitti & Siegelbaum, 2014; Meira et al., 2018). CA2 neurons in the dorsal hippocampus (dCA2) primarily send their axonal projections within the hippocampal formation (Cui, Gerfen, & Young, 2013; Hitti & Siegelbaum, 2014), including ventral CA1 (vCA1), which itself has been implicated in social memory (Okuyama, Kitamura, Roy, Itohara, & Tonegawa, 2016). Multiple lines of investigation suggest that projections from dCA2 to vCA1 are critical for social recognition (Meira et al., 2018; Raam, McAvoy, Besnard, Veenema, & Sahay, 2017), but how CA2 activity modulates CA1 is incompletely understood.

Activation of OXTRs in CA2 is known to increase neuronal excitability and cause local pyramidal neurons to enter into a burst firing mode (Tirko et al., 2018), as well as potentiate synaptic transmission onto CA2 neurons (Pagani et al., 2015). To bridge our understanding of oxytocinergic modulation at the cellular level with observations made in behavioral experiments, we have studied the circuit consequences of oxytocin’s actions in CA2, focusing on which CA2 cell-types are modulated, how the cellular response evolves over time and how CA2 activity propagates to other regions. Our experiments reveal a series of unexpected sequelae that occur in CA2 pyramidal cells upon OXTR activation to produce a persistent change in firing mode that occurs over a behaviorally relevant timescale. This persistent modulation is specific to CA2 pyramidal cells, which, in turn, strongly target *stratum oriens*-resident interneurons in CA1. Release of endogenous oxytocin into the hippocampus elicits sustained plasticity in CA1 pyramidal cells, a potential circuit mechanism to translate transient oxytocin release into persistent modifications in hippocampal activity as may occur during a social encounter.

## Results

Stimulation of hippocampal OXTRs excites both pyramidal cells and parvalbumin-expressing (PV+) interneurons via closure of the “M-current” or I_M_, which is mediated by potassium-fluxing KCNQ channels (Tirko et al., 2018). However, responses in the two cell-types show remarkably different time-courses (**Fig. 1a, b**). In these experiments, the specific OXTR agonist Thr^4^-Gly^7^-Oxytocin (TGOT, 400 nM) was applied to acute hippocampal slices from adult mice of either sex while whole-cell recordings were made from CA2 cells. TGOT was applied to each slice only once, to avoid receptor internalization as a confounding factor (Busnelli et al., 2012; Gimpl & Fahrenholz, 2001; M. P. Smith et al., 2006). CA2 pyramidal neurons were identified on the basis of their characteristic electrophysiological properties (Chevaleyre & Siegelbaum, 2010; Dudek et al., 2016; Tirko et al., 2018; Zhao, Choi, Obrietan, & Dudek, 2007) and transgenically labeled PV+ interneurons in CA1 and CA2 were visually identified by tdTomato expression (**Fig. 1a inset**). Both cell-types depolarized in response to TGOT stimulation within 2 min (average time to depolarization: 0.74 ± 0.22 min (PYR) vs. 0.54 ± 0.15 min (PV); **Fig. 1c**), but only pyramidal cells mounted a response that lasted tens of minutes (as shown in **Figs. 1b, d, e, f**). As a population, CA2 pyramidal cells exhibited a significantly longer lasting burst response to TGOT stimulation than PV+ interneurons (median burst duration: 18.6 min (PYR) vs. 7.6 min (PV), **Fig. 1d**). In contrast, PV+ cells consistently showed a large and transient increase in firing (average change in peak firing rate: 20.6 ± 3.5 Hz, n = 12) that roughly matched the duration of TGOT application. Of 12 PV+ cells recorded, all 12 depolarized in response to TGOT application (average peak depolarization: 14.3 ± 1.3 mV, n = 12), suggesting an absence of sub-type specificity in TGOT sensitivity. To define which subclasses of PV+ interneurons were included in our data set, we generated 2D morphological reconstructions after each recording and classified neurons on the basis on their axonal arborizations (**Fig. 1g**). The response to TGOT was indistinguishable between the two observed PV+ subtypes: the mean depolarization was 10.6±0.9 mV in bistratified neurons (5/9 reconstructed cells) and 9.8±1.6 mV in perisomatic targeting neurons (4/9 reconstructed cells)(p = 0.64, unpaired *t*-test).

**Figure 1.**
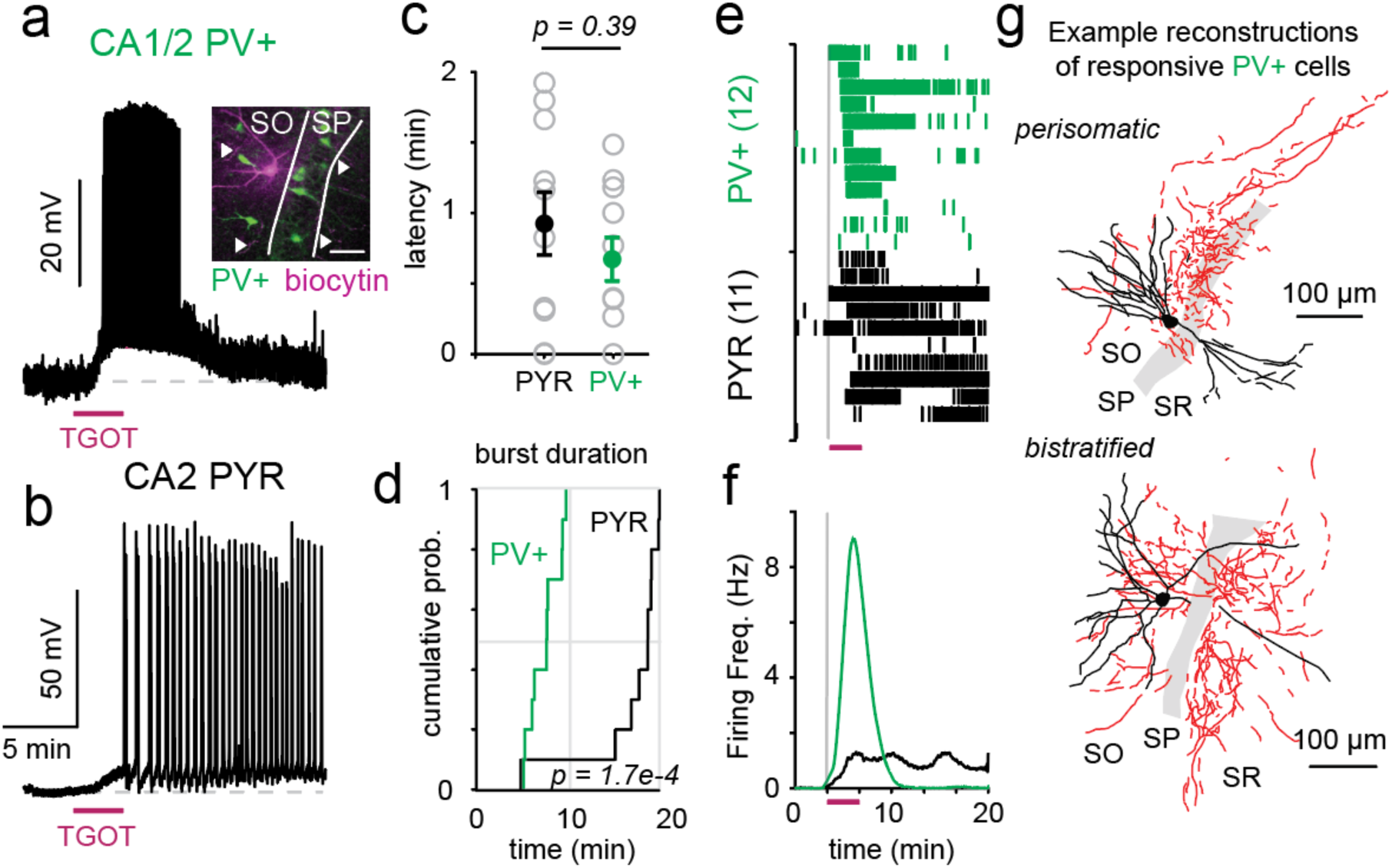
Excitatory and inhibitory hippocampal neurons show cell-type specific responses to OXTR stimulation. PV+ interneurons in CA1 and CA2 (a) and CA2 pyramidal cells (b) are depolarized by OXTR stimulation (TGOT, 400 nM, bath application). PV+ interneurons were targeted for recording using a transgenic mouse line (PV-*ires*-Cre X Ai9). Inset shows an example recorded cell filled with biocytin (streptavidin labeling in purple) and PV-expressing cells in green). The pyramidal cell layer, *stratum pyramidale* (SP), is outlined in white to distinguish it from the *stratum oriens* (SO) layer. Arrows point to axonal arborizations. Scale bar = 50 μm. Response latencies were comparable between cell-types; p = 0.39, unpaired *t*-test (c), while burst durations sharply differed, p = 1.7e-4, Kolmogorov-Smirnov test (d). Raster plots for CA2 neurons are shown in (e), which are collapsed to reflect instantaneous firing frequency in (f). Numbers in parentheses indicate group sizes for data in c – f. Example 2D morphological reconstructions of PV+ neurons are shown in (g). The soma and dendrites are shown in black, while the axon is in red. The SP is demarcated in gray. SR refers to the *stratum radiatum*. PYR group data from 11 cells / 10 mice. PV group data from 12 cells / 5 mice. 1 supplemental figure.

In the prefrontal cortex, a specific class of somatostatin-expressing (SST+) interneurons express the OXTR and are implicated in social-sexual behavior (Nakajima et al., 2014). To test for the recruitment of hippocampal SST+ interneurons by OXTR stimulation, we recorded from fluorescently labeled SST-expressing CA2 interneurons in a transgenic mouse line (SST-Cre x Ai9). The TGOT responses were generally small in this group (2.5 ± 0.8 mV, n=9), though 3 of 9 SST+ cells did display burst firing (**Fig. 1 – Supp. 1**). The magnitude of TGOT-induced depolarization was variable even within a defined SST+ subclass: anatomically confirmed OLM (*oriens-lacunosum moleculare*) interneurons (**Fig. 1 – Supp. 1**).

In contrast to the PV+ interneuron response, CA2 pyramidal cell responses were highly variable and often long outlasted the stimulus (**Fig. 1**). We next considered what mechanisms might underlie the cell type-specific persistence of this response to OXTR stimulation. Because TGOT responses in CA2 pyramidal cells are long-lasting even in the presence of excitatory synaptic blockers (Tirko et al., 2018), we focused on intracellular, not synaptic, signaling mechanisms. We first asked whether or not bursting activity was perpetuated via a “latch” mechanism, whereby once the cell started spiking, it entered a self-perpetuating bursting state. To test this, we forced CA2 pyramidal cells to burst repetitively by injecting ramps of depolarizing current (reaching ∼300 pA over 6 s; **Fig. 2a**). This stimulus caused CA2 PYRs to fire ∼35 action potentials, well within the range of what is observed upon OXTR stimulation. Despite multiple forced burst events, the CA2 PYR membrane potential (V_m_) remained unchanged without active current injection (pre vs post: -64 ± 1.5 mV vs. -65.2 ± 1.4 mV, p = 0.44, paired *t*-test; **Fig. 2a**). Self-perpetuating burst firing was never observed.

**Figure 2.**
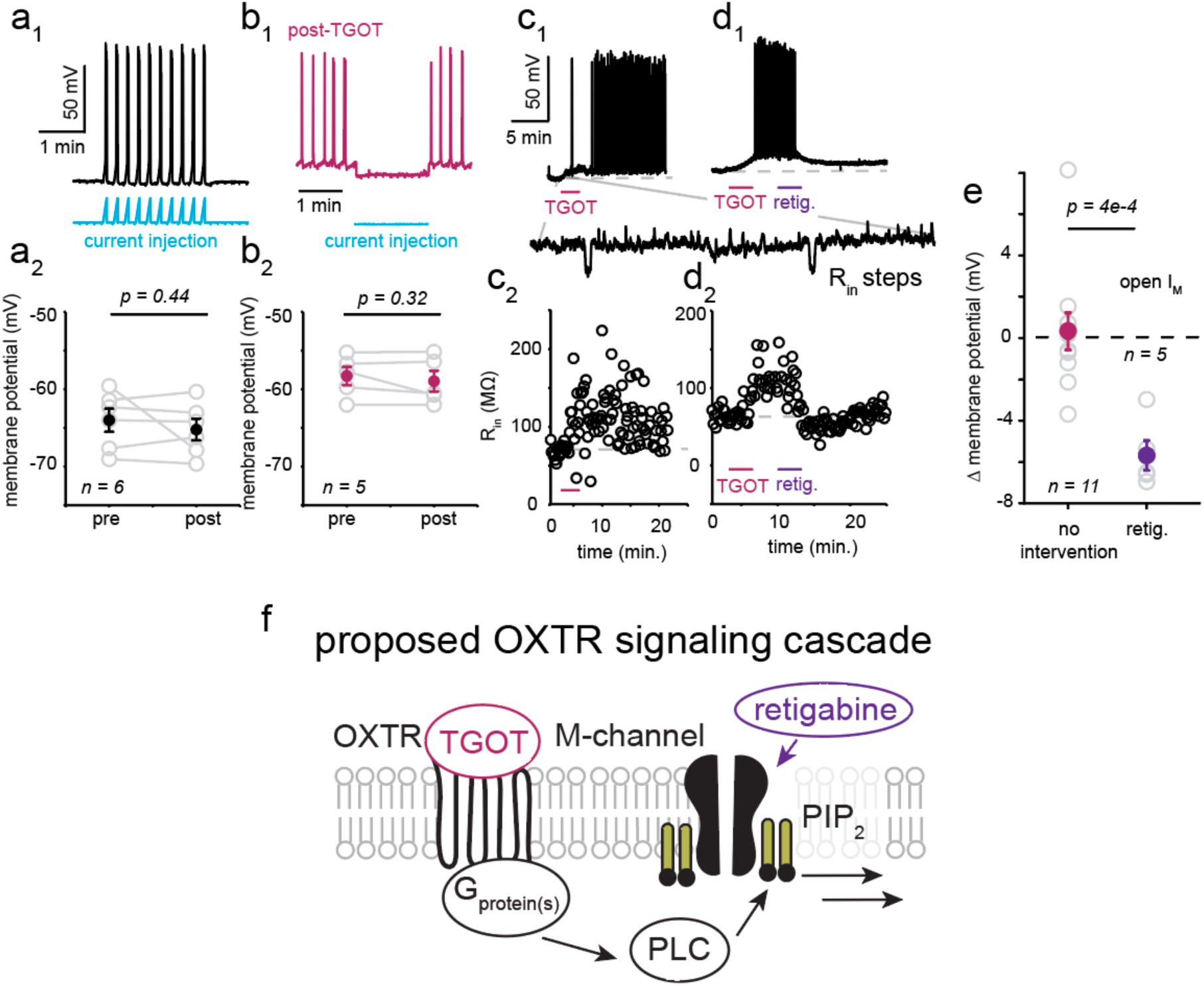
Sustained M-current inhibition is responsible for persistent OXTR responses. When forced to fire bursts of action potentials by a ramping current injection at the soma (example cell in a1), the membrane potential of CA2 pyramidal cells is unchanged (a2; p = 0.44, paired *t*-test; n = 6 cells / 5 mice). Imposition of a hyperpolarizing current for 200 seconds did not return the TGOT-excited cell to its baseline membrane potential (b; n = 5 cells / 4 mice). Input resistance and membrane potential remain elevated following TGOT application in CA2 pyramidal cells (c). Application of the KCNQ channel opener, retigabine (100 μM, retig.), repolarized the TGOT-excited CA2 PYRs and returned cellular input resistance to baseline values (d). Group data summarized in (e). Control group data comes from 11 cells / 8 mice. Retigabine group data from 5 cells / 3 mice. All error bars reflect the s.e.m. Schematic of known signaling machinery downstream of the OXTR in CA2 cells (modified from (Tirko et al., 2018), (f)). One supplemental figure.

In a complementary series of experiments, we asked whether the persistent response was dependent on a continually depolarized membrane potential. After inducing burst firing and depolarization by TGOT application, we held the cell at a hyperpolarized membrane potential (hyperpolarizing the cell by ∼8 mV, to near baseline potential, for 200 s; **Fig. 2b**). To our surprise, this sustained hyperpolarization was unable to trigger a return to the cell’s baseline state (pre vs post: -58.3 ± 1.2 mV vs. -59 ± 1.3 mV, p = 0.32, paired *t*-test). After cessation of the hyperpolarizing current, cells went right back to a depolarized membrane potential and burst firing (**Fig. 2b**). Thus, depolarizing the cell was not enough to induce continual burst firing (**Fig. 2a**) and hyperpolarizing the cell during TGOT-induced bursting was not sufficient to stop previously triggered activity.

These results prompted us to consider possible biochemical underpinnings of the persistent activity. In response to OXTR stimulation, CA2 PYR cell input resistance increased and remained elevated for the duration of the voltage response (**Fig. 2c, Fig. 2 – Supp. 1**). Previously, we pharmacologically demonstrated that OXTR stimulation increased input resistance due to closure of the M-channel (Tirko et al., 2018). This observation led us to suspect that sustained channel inhibition might be responsible for the long-lasting depolarization and burst firing, even though swifter recovery from inhibition has been found in other neurons, like sympathetic ganglion cells (Suh & Hille, 2002). To test explicitly for sustained I_M_ inhibition, we applied the M-current opener, retigabine (100 μM), a few minutes after TGOT removal to determine if this could reverse TGOT-induced excitation (**Fig. 2d**). Input resistance (R_in_), an indirect assay of M-current conductance, was sampled every 10 s with a small hyperpolarizing current step (magnified trace, **Fig. 2c, d**). Application of retigabine promptly reversed the depolarization, repetitive firing and change in R_in_ induced by TGOT (mean change in Vm: -5.68±0.7 mV, n = 5; **Fig. 2d, e**). Simply waiting 10 minutes after TGOT application did not result in a comparable repolarization (mean change in Vm: 0.31±0.9 mV, n = 11, p = 0.74; **Fig. 2c**). The ability of retigabine to reverse TGOT’s long-lasting effects is consistent with OXTR stimulation causing a *sustained* inhibition of I_M_.

M-current inhibition is most often driven by depletion of phosphatidylinositol-4,5- biphosphate (PIP_2_) from the plasma membrane by phospholipase C (PLC, **Fig. 2f**) as KCNQ channels require PIP_2_ to open (Suh & Hille, 2002; Zhang et al., 2003). We have previously reported that pre-treatment with U73122, commonly employed as a PLC inhibitor, prevents TGOT-mediated excitation (Tirko et al., 2018). We next tested whether U73122 treatment was also able to reverse OXTR-driven depolarization. Unlike retigabine, U73122 was unable to return CA2 PYRs to their resting Vm or input resistance (mean change in Vm following U73122 treatment: 0.31±0.9 mV, n = 11; p = 0.37). These data are consistent with sustained PLC activation being dispensable for sustained M-current inhibition.

Classically, M-current inhibition is induced following activation of Gαq, which directly interacts with PLC to degrade membrane PIP_2_. The OXTR is capable of coupling to both Gq and Gi α proteins (Gravati et al., 2010; Hoare et al., 1999; Rimoldi et al., 2003; Strakova & Soloff, 1997; Zhou, Lutz, Steffens, Korth, & Wieland, 2007), raising the possibility that non-canonical G-protein signaling might account for the prolonged inhibition of the M-channel. To explore this idea, we asked 1) whether other neuromodulatory receptors, which signal though Gαq alone, were sufficient to produce sustained bursting in CA2 cells and 2) what G-proteins are required for TGOT-induced burst firing in CA2 pyramidal cells.

The AVP1bR subtype of vasopressin receptor is expressed in CA2 pyramids and is thought to be Gαq-coupled (Pagani et al., 2015), prompting us to compare the voltage response to oxytocin stimulation to that of vasopressin. Application of arginine vasopressin (AVP, 1 μM), depolarized CA2 PYRs and elicited bursts of action potentials that were similar in mean frequency (TGOT v AVP: 24.3±5.6 vs. 12.2±2.8 Hz; p = 0.20, unpaired *t*-test) and duration (TGOT v AVP: 14.4±1.6 vs. 18.3±2.4 min; p = 0.34, unpaired *t*-test) to those elicited by TGOT (**Fig. 3 – Supp. 1**). The similarity between AVP and TGOT responses suggested that signaling through Gαq proteins alone, via the AVP1bR, was capable of producing a long-lasting depolarization. While AVP is capable of signaling through both the OXTR and the AVP1bR (Song & Albers, 2018), application of AVP is capable of producing sustained depolarization in OXTR KO animals (**Fig. 3 – Supp. 2**).

Both TGOT and AVP elicit bursting firing in CA2 pyramidal cells, but only TGOT caused highly variable responses. We hypothesized that some of this variability might be due to the activation of multiple G-proteins downstream of the OXTR. To first test whether the OXTR signaled via Gα_q_, whose activation directly stimulates PLC, we measured the TGOT response in slices pre-treated with the specific Gα_q_ -family inhibitor FR900359 (FR), (Schrage et al., 2015). FR pre-treatment (1 μM, 1 h) blocked the TGOT response (control v. drug-treated: 5.6±0.8 mV (TGOT) v. 0.9±0.5 mV (TGOT+FR); p = 0.001; unpaired *t*-test), along with that of a positive control, the depolarization induced by the cholinergic agonist carbachol (control v. drug-treated: CCh, 9.6±2.6 mV (CCh) vs. 0.9±0.8 mV (TGOT+FR); p = 0.009, unpaired *t*-test; **Fig. 3a-c**). Carbachol is known to induce burst firing in CA2 pyramidal cells by signaling through M1 and M3 muscarinic receptors that are classically Gα_q_-coupled (Robert et al., 2020). FR responsiveness implies the involvement of a Gα_q_ protein in both the TGOT and CCh response, as would be expected from the literature.

**Figure 3.**
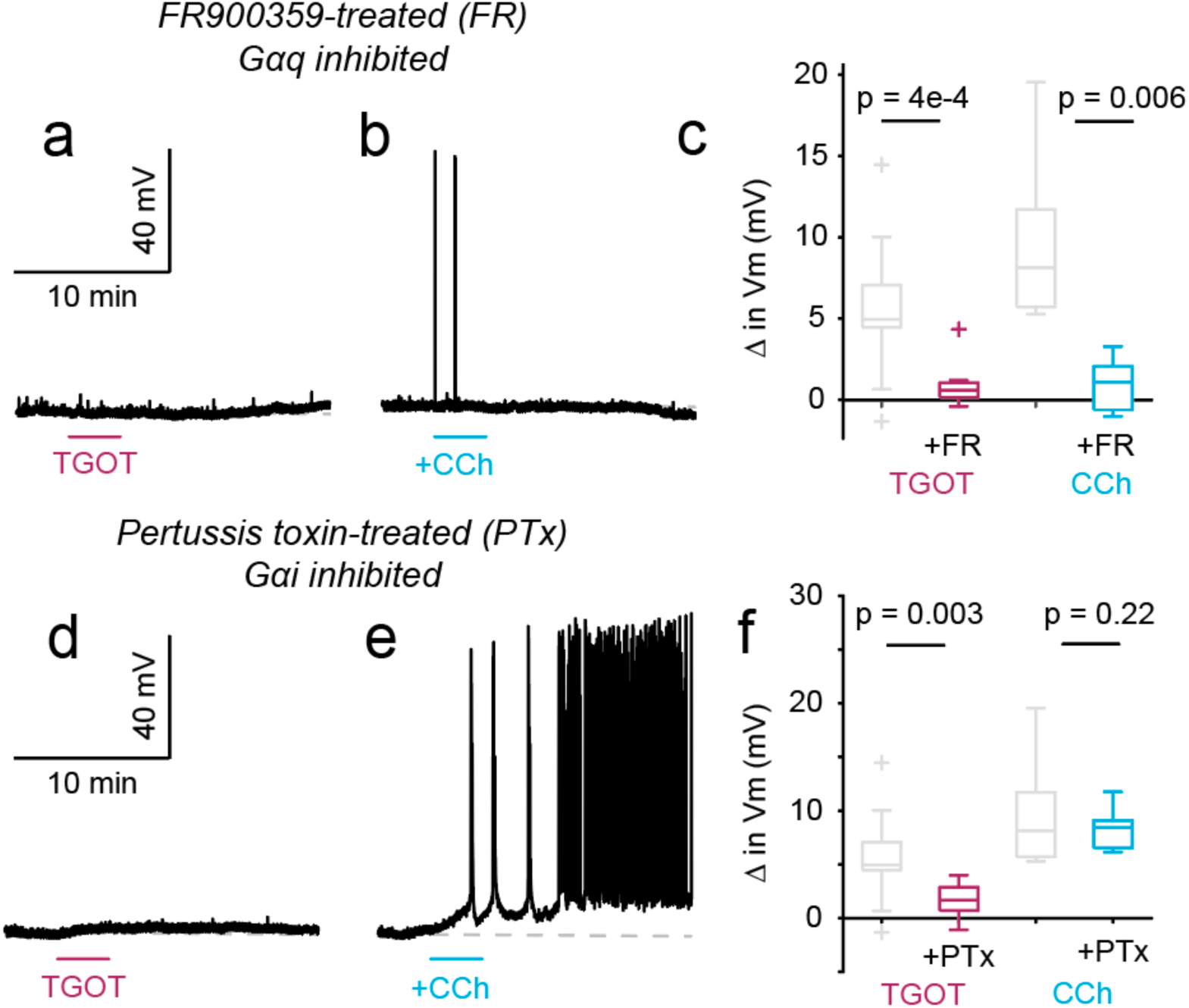
Multiple modulators elicit burst firing in CA2 PYRs, via different G-protein signaling mechanisms. Pre-treatment with FR900359 blocks the response to OXTR (a) and acetylcholine receptor (b) activation. Group data summarized in (c). Pre-treatment with pertussis toxin blocked TGOT-induced depolarization in CA2 PYRs (d), but not the response to carbachol, CCh, (e). Group data summarized in (f) as the mean change in membrane potential after drug treatment. Results of one-way ANOVAs (*p = 0.001* and *p = 0.004* for TGOT and CCh comparisons, respectively) prompted us to make pairwise comparisons between groups, which are reported in text with Tukey-Kramer correction for multiple comparisons. Group sizes are as follows: TGOT only (18 cells / 12 mice); TGOT+FR (9 cells / 7 mice); CCh only (5 cells / 2 mice); CCh+FR (5 cells / 2 mice); TGOT+PTx (9 cells / 5 mice); CCh+PTx (6 cells / 3 mice). 4 supplemental figures.

In an independent series of experiments, we pre-treated mice with the specific Gα_i_ inhibitor pertussis toxin (PTx; intraventricular injection, 24-72 hours before preparing slices). The long pre-treatment with PTx did not alter the electrical properties of CA2 PYRs or the frequency of synaptic input (**Fig. 3 - Supp. 3**). PTx pre-treatment did, however, blunt the depolarization usually caused by TGOT (control v. drug-treated: 5.6±0.8 mV (TGOT) v. 1.7±0.5 mV (TGOT+PTx) p = 0.02; unpaired *t*-test), but not that caused by CCh (control v. drug-treated: 9.6±2.6 (CCh) v. 8.4±0.8 (CCh+PTx) mV; p = 0.64; unpaired *t*-test; **Fig. 3d-f**). Like FR, PTx pre-treatment also inhibited the TGOT- induced increase in input resistance (**Fig. 3 - Supp. 4**). This PTx sensitivity was specific to pyramidal cells; CA2 PV+ interneurons treated with PTx still responded to TGOT (mean depolarization: 8.8 ± 1.8 mV, n = 3). Furthermore, the PTx sensitivity did not extend to CCh responsiveness (**Fig. 3e**), indicating that involvement of Gα_i_ is not a general prerequisite for persistent bursting.

Sensitivity to both PTx- and FR-treatment suggest that OXTR-induced burst firing in CA2 PYRs requires the activity of G_q_ and G_i_ α proteins. While PTx-sensitive proteins are capable of signaling to certain PLC isoforms via βγ-proteins (Camps et al., 1992; Katz, Wu, & Simon, 1992), their activation is not typically known to cause M-current inhibition. It is equally surprising to find a GPCR whose neuronal signaling requires activation of *both* Gα_q_ and Gα_i_, although there is precedent for such joint dependence in immune cells (Shi et al., 2007).

Independent of receptor type or G-protein(s) involved, burst firing of CA2 PYRs appears a widespread phenomenon across modulatory systems. To begin to understand the functional significance of CA2 burst firing, we considered how it might propagate to downstream regions. While CA2 PYRs project to numerous areas of the brain, most axonal fibers converge on the neighboring hippocampal sub-region CA1 (Cui et al., 2013; Hitti & Siegelbaum, 2014). To visualize CA2 PYR projections there, we delivered a Cre-dependent virus encoding ChETA-YFP into the CA2 regions of Amigo2-Cre mice and quantified YFP signal density (**Fig. 4a**). Consistent with reports from other groups (Dudek et al., 2016; Hitti & Siegelbaum, 2014; Kohara et al., 2014; Tamamaki, Abe, & Nojyo, 1988), we observed that axonal projections were densely concentrated in the interneuron-filled *stratum oriens* (SO) layer of CA1, with substantial projections to the *stratum radiatum* (SR) as well. Projections to SO showed significantly more labeling than the *stratum pyramidale* (SP; 24±2.8 (SO) vs. 8.2±1.2 (SP): p = 3.6e-5; paired *t*- test). Basal CA1 PYR cell dendrites and local interneurons are positioned to be targeted by CA2 axons in the SO layer and direct synaptic connections between CA2 PYRs and CA1 PYRs have been reported previously (Chevaleyre & Siegelbaum, 2010; Hitti & Siegelbaum, 2014; Kohara et al., 2014). To evaluate the effect of hippocampal oxytocin on spontaneous activity in CA1, we recorded from pyramidal cells in current clamp during TGOT bath application, expecting to see synaptically-propagated EPSPs driven by CA2 PYR burst firing (as in example 1, **Fig. 4c**). To our surprise, however, we observed no change in EPSP frequency across the set of recordings (average change in EPSP frequency: 1±1.3 Hz; p = 0.44, one-sample *t*-test; **Fig. 4c,d**). This absence of TGOT-stimulated CA2 PYR drive onto excitatory CA1 cells was independent of distance from CA2 and CA1, and whether or not the CA1 PYR was in the deep or superficial pyramidal layer (data not shown). To understand why we did not observe significant synaptic excitation onto CA1 PYRs during TGOT presentation, we revisited CA2 PYR cell anatomy. As CA2 PYR axons most strongly innervate regions rich in interneurons (**Fig. 4a, b**), we sought to determine the relative strength of CA2 PYR cell synapses onto excitatory and inhibitory cells in the CA1 SO. In these experiments, we made serial recordings from neighboring CA1 PYR and SO interneuron “pairs”, while optogenetically stimulating CA2 PYR cell fibers and keeping the intensity of light stimulation the same. Consistently, SO-resident interneurons received significantly stronger CA2 input than nearby PYRs (average EPSP amplitude, SO v. PYR: 9.8±1.9 vs. 2.2±0.4 mV; p = 0.002, paired *t*-test, n = 10, **Fig. 4e, f**). Most interneurons fired action potentials in response to a single pulse of blue light, whereas CA1 PYRs were never brought to spike threshold with the same stimulus. As dCA2 projections to vCA1 have been shown to be critical for CA2’s role in social recognition (Meira et al., 2018; Raam et al., 2017), we also tested if this phenomenon would hold in ventral hippocampus, where, indeed, we observed a similar trend (average EPSP amplitude, SO v. PYR: 3.1±1.5 vs. 0.6±0.4 mV; p = 0.17, n = 3, paired *t*-test). We also considered the possibility that the subset of CA2 PYRs that express the OXTR (OXTR+) might show a different pattern of SO innervation than the cell population as a whole. To test this explicitly, we injected floxed ChETA-YFP into the CA2 sub-region of OXTR-*ires*-Cre animals and again recorded in CA1 “pairs” while optogenetically stimulating CA2 PYR cell fibers. As CA2 interneurons also express the OXTR and are likely to express virally delivered channelrhodopsin in experiments in the OXTR-*ires*-Cre line, we clamped the CA1 PYR cell voltage to -70 mV to isolate excitatory currents. Similar to what we observed when stimulating CA2 broadly, there was a trend for CA1 SO interneurons to receive stronger CA2 input than nearby PYRs (average EPSC magnitude, SO v. PYR: 207.1±77.8 vs. 74.9±24.2 pA, p = 0.11, n = 5, paired *t*-test).

**Figure 4.**
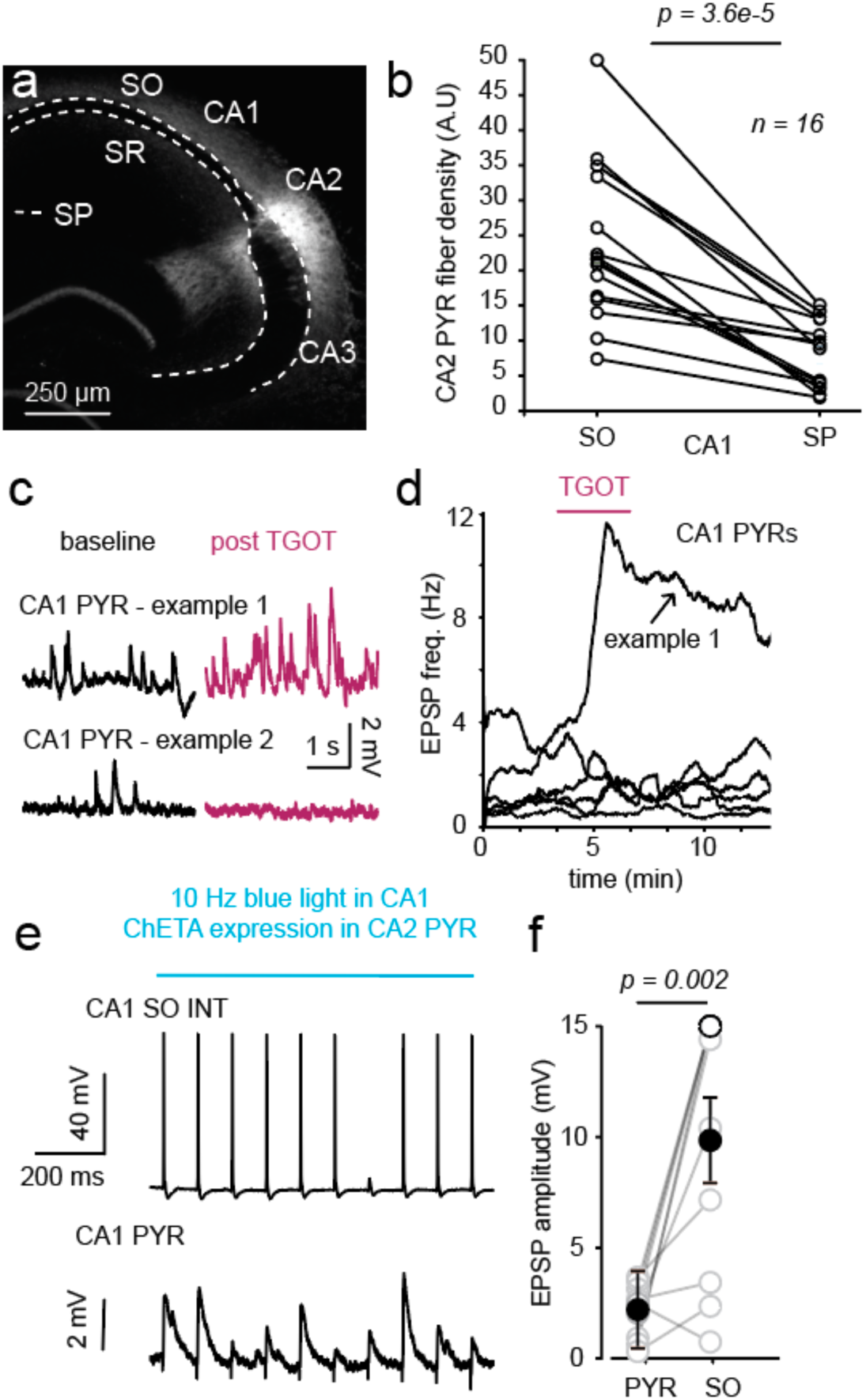
CA2 PYRs strongly innervate perisomatic CA1 SO interneurons. Viral expression of YFP in CA2 PYRs, driven by Amigo2-Cre, in an example slice in three sub- regions of CA1: *stratum oriens* (SO), *stratum pyramidale* (SP) and *stratum radiatum* (SR; a). Quantification of CA2 PYR cell fiber density in SP and SO across slices (b; n = 16 cells / 7 mice). Fiber density given in arbitrary units (A.U.). Example current clamp recordings from CA1 PYRs before and after TGOT application (c). Quantification of EPSP frequency in CA1 PYRs upon drug application (d; 6 cells / 4 mice). Postsynaptic responses recorded in an example CA1 SO interneuron (top) and pyramidal cell (bottom) upon blue light stimulation (10Hz, 1 s) of CA2 PYR fibers (e). Quantification of EPSP amplitude elicited by optogenetic stimulation of CA2 PYR fibers in dorsal CA1 (f; n = 10 cells / 7 mice). 1 supplemental figure.

Whenever we stimulated CA2 PYRs and recorded in CA1 SO interneurons, we observed a bimodal distribution in the interneuron population response. The majority of interneurons received a strong CA2 PYR cell input, while a minority displayed EPSPs on par with CA1 pyramids. To test if this bimodality was caused by differential targeting of interneuron subtypes by CA2 PYRs, we characterized SO interneurons on the basis of electrophysiological (input resistance and sag ratio as a proxy for I_h_ current) properties and axonal projection anatomy (**Fig. 4 - Supp. 1**). In this analysis, we identified multiple interneuron subclasses in our data set. In general, strongly targeted interneurons, (EPSP >5 mV) were characterized by significantly lower input resistance (R_in_, strongly v. weakly targeted: 77.7±14.7 v. 165±29.3 MΩ; p = 0.009, unpaired *t*-test) and less I_h_ (Ratio of sag current to steady state, evoked by a hyperpolarizing pulse, strongly v. weakly targeted: 0.13±0.05 v. 0.33±0.08, p = 0.04, unpaired *t*-test) than those that were weakly targeted (**Fig. 4 – Supp. 1**). The physiological properties of strongly targeted interneurons are consistent with features of fast-spiking PV+ interneurons, which may have perisomatic or bistratified axonal projections. Posthoc reconstructions of SO interneurons revealed that the interneurons receiving strong CA2 input classified as perisomatic, but not bistratified (**Fig. 4 – Supp. 1**). At least one additional class of interneurons, characterized by strong adaptation and large AHP, was also strongly innervated by CA2 PYRs.

We next asked how strong targeting of CA1 interneurons by CA2 PYRs might locally regulate evoked activity in CA1. First, we considered how acute stimulation of CA2 PYRs influences spike transmission between CA3 and CA1 PYRs, evoked via stimulation of Schaffer Collaterals (SC). To do so, we optogenetically mimicked CA2 burst firing by delivering light pulses at 20 Hz for 1 s to ChETA-bearing CA2 fibers in CA1 while simultaneously stimulating the SC (**Fig. 5 – Supp. 1**). Baseline spike probability in response to SC stimulation was established over 20 trials, before interleaving every other stimulus with delivery of the blue light. This burst-like stimulation of CA2 PYR fibers had no effect on CA3-CA1 spike transmission or EPSP amplitude (**Fig. 5 – Supp 1**), prompting us to ask if optogenetic release of oxytocin, which produces much more persistent burst activity in CA2 (Tirko et al., 2018), can influence CA3-CA1 transmission. Accordingly, we next subjected oxytocinergic fibers, which virally expressed ChETA-YFP and course through the hippocampus, to optogenetic stimulation with blue light pulses (30 Hz for 60 s). After obtaining a 5-minute baseline recording of SC-evoked EPSPs, we stimulated oxytocinergic fibers and observed a sustained, 2-fold increase in SC-evoked EPSP amplitude (post/pre: 2.04±0.4; p = 0.03, one-sample *t*-test; **Fig. 5a,b**). This potentiation was not seen when slices were pre-treated with the OXTR antagonist OTA (post/pre: 1.15±0.2; p = 0.392, one-sample *t*-test; **Fig. 5a, b**) or the GABA-A receptor blocker bicuculine (post/pre: 1.4±0.4; p = 0.393, one-sample *t*-test). As an indication of cell health and recording stability, we continuously monitored input resistance, which remained stable throughout the recording period (**Fig. 5 – Supp. 2**). The enhancement of evoked CA3-CA1 transmission was not simply due to direct synaptic modulation by oxytocin, insofar as TGOT did not affect the amplitude or dynamics of SC-evoked synaptic currents in CA1 pyramidal cells (**Fig. 5 – Supp. 3**). Consistent with a role for interneurons in this phenomenon, we observed a trend for the compound IPSP to enlarge upon oxytocinergic stimulation (Change in net IPSP: 1.07±0.5 mV; p = 0.07, one-sample *t*- test) that was not observed following OTA pre-treatment (0.59±0.5 mV; p = 0.36, one-sample *t*-test; **Fig. 5c**). Also consistent with activation of interneurons, the SC-evoked EPSP narrowed following optogenetic stimulation (change in EPSP width: -0.99±0.4 mV; p = 0.04, one-sample *t*-test), while OTA pre-treated slices actually showed a broadening of the PSP waveform (0.77±0.19 mV; p = 0.03, one-sample *t*-test; **Fig. 5d**).

**Figure 5.**
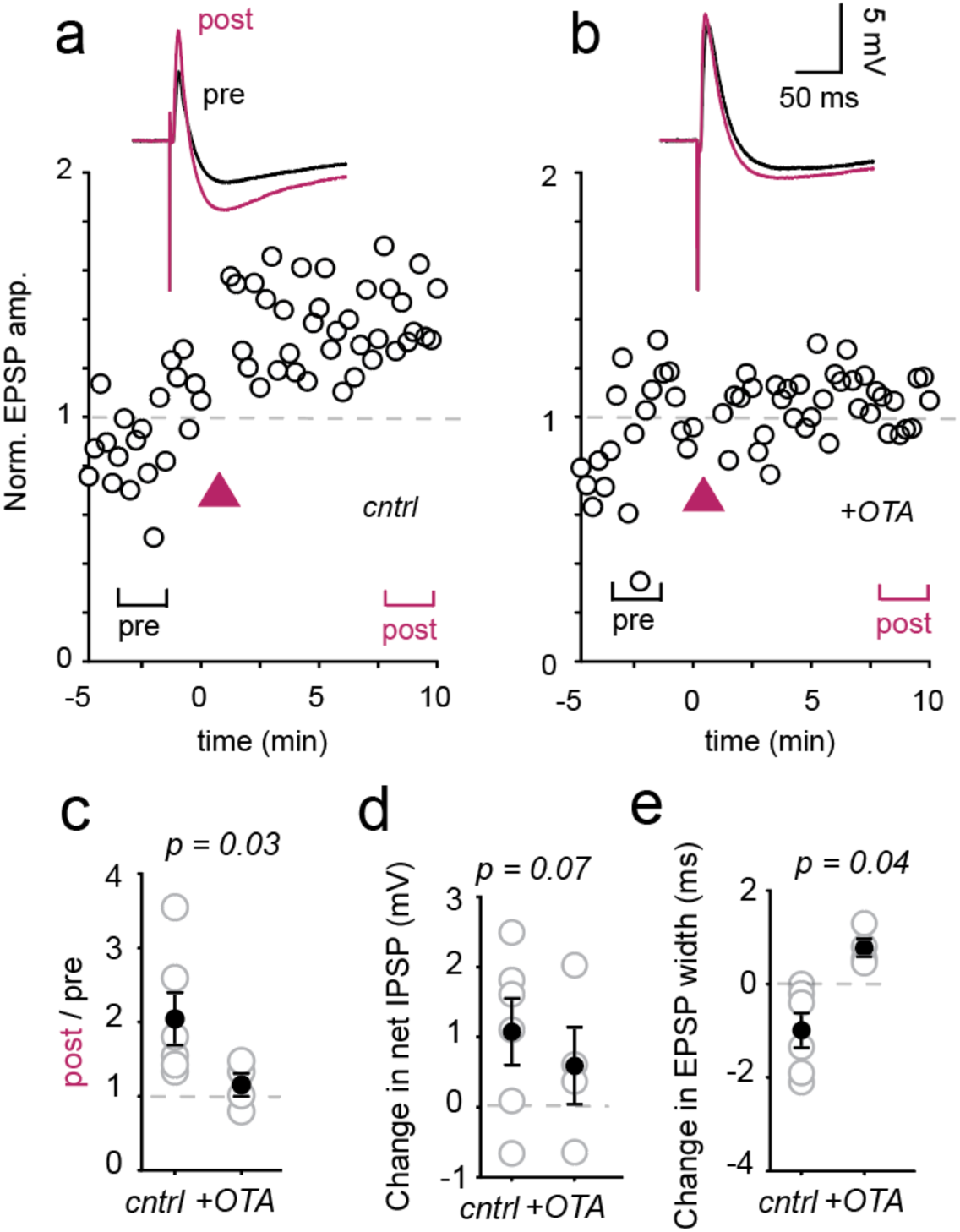
Oxytocinergic activation of the hippocampus shapes CA3-CA1 transmission. (a) SC evoked EPSPs before (pre) and after (post, magenta) blue light stimulation of oxytocinergic fibers. Time course of change in EPSP amplitude in example cell plotted below. (b) Results of the same protocol performed in the presence of OXTR blockade with the OXTR antagonist OTA (1 μM). (c) Quantification of change in EPSP amplitude from baseline (pre) to 10 minutes after stim (post). (d) Amplitude of the net IPSP recorded after stimulation subtracted by that measured at baseline. (e) EPSP width recorded after stimulation subtracted by that measured at baseline. P-values reflect results of a paired *t*-test. Control group data comes from 6 cells / 4 mice. OTA group data comes from 4 cells / 3 mice. 4 supplemental figures.

Oxytocin will depolarize both CA2 (Tirko et al., 2018) and CA3 pyramidal neurons (Lin, Chen, Huang, Nishimori, & Hsu, 2017), raising the possibility that depolarization of CA3 pyramidal cells might account for the enhanced EPSP. To disambiguate the contribution of CA3 neurons from CA2 neurons, we recorded the amplitude of the fiber volley in CA3 pyramidal cell axons upon optogenetic release of endogenous oxytocin; the amplitude of the SC fiber volley was unchanged (**Fig. 5 – Supp. 4**). In sum, CA1 pyramidal cell input resistance (**Fig. 5 – Supp. 2**), evoked CA3-CA1 EPSC amplitude (**Fig. 5 – Supp. 3**) and CA3 axonal excitability (**Fig. 5 – Supp. 3**) were all unchanged in response to oxytocin release, lending support for an underlying circuit mechanism for the enhancement and narrowing of the evoked EPSP.

## Discussion

The association of hippocampal sub-region CA2 with social behavior is now well-established (Hitti & Siegelbaum, 2014; Meira et al., 2018; A. S. Smith, Williams Avram, Cymerblit-Sabba, Song, & Young, 2016), as is the need for OXTR expression in CA2 pyramidal cells during social recognition tasks (Lin et al., 2018; Raam et al., 2017). But how oxytocin signaling in CA2 promotes formation of a social memory remains a major open question. In an effort to understand how oxytocin modulates the underpinnings of social information processing, we studied the circuit consequences of oxytocin signaling in hippocampal area CA2. We report that while both CA2 pyramidal cells and CA1/CA2 interneurons respond to OXTR stimulation (Owen et al., 2013; Tirko et al., 2018), burst firing and increased input resistance persist specifically in CA2 excitatory cells. This sustained activity in CA2 PYRs is accompanied by elevated input resistance and is reversed by application of the M-current opener retigabine, indicating that it is due to prolonged inhibition of the M-current.

How the M-current is inhibited for tens of minutes at a time remains an open question. Gene expression analysis from CA2 pyramidal cells provides some possible answers (Cembrowski, Wang, Sugino, Shields, & Spruston, 2016). PI4K, the rate-limiting enzyme in PIP_2_ re-synthesis (Suh & Hille, 2002), is expressed at lower levels in CA2 relative to CA1 and CA3 pyramidal cells (Cembrowski et al., 2016). Alternatively, slow and long-lasting inhibition of M-current can also be induced by receptor tyrosine kinases that have been activated by Gα_i_ proteins (Gamper, Stockand, & Shapiro, 2003; Jia et al., 2007).

The complete block of OXTR signaling in CA2 pyramidal cells after FR treatment leads us to believe the Gα_q_-pathway is the dominant signaling mechanism. Surprisingly we found an additional effect of blocking G_i_ signaling; only a small depolarization remains in PTx-treated neurons. The unique Gα_q_/ Gα_i_ requirement for OXTR signaling in CA2 pyramids may play a role in the unusual temporal dynamics of the burst response. Synergistic signaling between G_αq_ and G_αi_ has been reported previously (Philip, Kadamur, Silos, Woodson, & Ross, 2010; Pierce, Mehrotra, Mustoe, French, & Murray, 2019; Rebres et al., 2011; Shah et al., 1999; Zhu & Birnbaumer, 1996) and might lead to persistent calcium signaling or PLC activation, explaining the long-term channel inhibition in terms of supra-additive PLC activation. Likewise, dual G-proteins are deployed downstream of beta-adrenergic subtype 2 receptors, which signal through both G_αs_ and G_αi_, in series to produce a transiently enhanced contraction rate in cardiac myocytes (Devic, Xiang, Gould, & Kobilka, 2001; Strohman et al., 2019). Signaling through multiple G-proteins may allow for the modulation of multiple classes of ion channel by multiple GPCRs as has been reported in CA2 pyramidal cells upon cholinergic stimulation (Robert et al., 2020).

To place our findings in broader physiological context, we considered how burst activity in CA2 neurons might impact downstream regions. We focused our efforts on the neighboring region CA1, wherein pyramidal cells do not express the OXTR, and most CA2 pyramidal cell axons converge (Cui et al., 2013; Hitti & Siegelbaum, 2014; Tirko et al., 2018). In addition to targeting CA1 pyramidal (Chevaleyre & Siegelbaum, 2010; Hitti & Siegelbaum, 2014; Kohara et al., 2014) and giant radiatum cells (Nasrallah et al., 2019), CA2 neurons form strong synaptic connections onto CA1 oriens-residing interneurons, including those innervating the pyramidal cell layer. While this E-I projection does not appear to regulate spontaneous CA1 pyramidal cell activity, it does refine evoked CA3-CA1 synaptic transmission and likely has implications for the timing and efficacy of spike propagation in CA1 pyramidal cells via feed-forward inhibition (Boehringer et al., 2017; Nasrallah et al., 2019; Owen et al., 2013; Pouille & Scanziani, 2001). The anatomical bias for strong CA2 input into the SO layer was observed in both dorsal and ventral hippocampal slices, which is particularly intriguing in light of recent work showing that PV+ ventral CA1 interneurons increase their activity in the presence of novel social stimuli and their loss impairs social recognition (Deng, Gu, Sui, Guo, & Liang, 2019). It is important to note that a minority of CA1 pyramidal cells, across different data sets, did show an increase in excitatory input upon TGOT application to the slice (**Fig. 4; Fig. 5 – Supp. 3**). These cells may represent a functionally distinct CA1 pyramidal subclass that is uniquely targeted by CA2.

Evoked firing of oxytocinergic fibers potentiated the excitatory component of evoked synaptic transmission at the CA3-CA1 synapse. The compound EPSP was both larger and briefer, an overall increase in strength and temporal precision lasting for the duration of the recording. We posit that in causing this long-lasting enhancement of CA3-CA1 transmission, endogenous release of oxytocin operates somewhat like flipping a record switch during a social encounter. The sustained response to transient oxytocin exposure might thus extend over a behaviorally relevant time scale to amplify and sharpen hippocampal neurotransmission, supporting the encoding of relevant spatial, sensory and social cues. Consistent with the idea, dCA2 to vCA1 transmission is critical during the encoding and consolidation phases of a 15-minute social encounter (Meira et al., 2018), as if dCA2 were activated early on, presumably by oxytocin, and continues to signal for tens of minutes. Such persistence of action appears to be a feature of oxytocin signaling across the brain. Acute oxytocin refines auditory cortical spike timing in response to pup calls for at least 2 hours after drug delivery (Marlin, Mitre, D’Amour J, Chao, & Froemke, 2015). Similarly, a single dose of intranasal oxytocin can improve social behavior in an animal model of autism spectrum disorder for 2-3 hours following application (Penagarikano et al., 2015). Thus, the influence of oxytocin *in vivo* far outlasts the likely duration of the ligand-surface receptor duration, consistent with the kind of temporal expansion reported here.

It is of considerable interest to decipher how the excitatory component of the compound CA3-CA1 PSP is refined and enhanced by oxytocin. The observations that 1) CA1 pyramidal cell membrane potential and input resistance are unchanged upon TGOT exposure or endogenous oxytocin release (**Fig. 5 – Supp. 2**) and 2) CA3 pyramidal cell fiber volley (**Fig. 5 – Supp. 2**) is unaffected by endogenous oxytocin release weigh against modulation of CA1 or CA3 cell excitability accounting for the potentiation. Similarly, because TGOT did not modulate the amplitude or synaptic dynamics of SC-evoked EPSCs, we do not think that oxytocin directly affects CA3 presynaptic release (**Fig. 5 – Supp. 3**). We propose that a circuit mechanism, perhaps involving interneuron modulation, may underlie the observed increase in EPSP amplitude. Modulation of inhibitory output is implied by the narrowing of the compound PSP and may have been predicted by increased excitability of CA1 interneurons (caused directly by their OXTR stimulation, or indirectly by excitatory synaptic drive coming from CA2 pyramidal neurons). Of these two possibilities, we regard CA2-CA1 interneuron drive to be the weightier contributor to the increased IPSP, which was observed 10+ minutes after light stimulus, well after the direct effect of oxytocin has subsided in interneurons.

We have previously reported that TGOT application acutely *reduces* evoked inhibition in the CA1 region of juvenile rats via activity-dependent depression of inhibitory synaptic output (Owen et al., 2013). Here we observed a similar decrease in feed-forward inhibition immediately following TGOT exposure in CA1 pyramidal cells of adult mice (**Fig. 5 – Supp. 3**). It remains open whether these acute responses are linked via disinhibition to the induction of sustained excitatory synaptic enhancement (Fig. 5a,b), or to the later strengthening of disynaptic inhibition (Fig. 5c), which may be secondarily linked to EPSP potentiation (D’Amour & Froemke, 2015).

Observing an OXTR-dependent effect on synaptic plasticity of inputs to CA1 offers fresh perspective on published behavioral findings that we now briefly discuss. A mouse’s ability to recognize familiar mice is impaired by toxin block of CA2 pyramidal cell output (Hitti and Siegelbaum 2014) and by genetic silencing of CA2 pyramidal cell axon terminals in ventral CA1 (Meira et al., 2018; Raam et al., 2017). These elegant studies established the importance of CA2 output for social recognition memory (SRM) and are further complemented by experiments that discriminate between plasticity at CA2→CA1 PYR synapses *per se* (Tirko et al., 2018) and a heterosynaptic influence on plasticity on other synapses targeting CA1 PYR (**Fig. 5**). The indirect promotion of synaptic potentiation could provide modulatory enhancement of the storage of incoming information via more classical pathways (e.g. CA3→CA1), switching on recording in dorsal CA1 to support object recognition and in ventral CA1 to promote social memorization, as functionally separated with optogenetics (Raam et al., 2017).

Potentiation of CA2 input synapses was not evident in our experiments, although we note that application of TGOT in hippocampal slices can promote NMDAR-dependent LTP (Pagani et al. 2015) and deletion of OXTR impairs potentiation onto CA2 PYRs (Lin et al., 2018). A critical observation is that NMDAR expression in CA2 PYR, while required for OXTR-induced potentiation of EPSCs (Pagani et al. 2015), can be deleted without affecting aggressive behavior or short-term social memory (Williams Avram et al., 2019). This, taken together with our observations on synaptic plasticity (**Fig. 5**), suggests that the most critical form of plasticity might occur downstream of burst activity in CA2 pyramidal cells and not in those neurons themselves.

Perhaps the biggest open question is when peptides like oxytocin and vasopressin, are released during behavior. While the peptidergic tone of the hippocampus during behavior is uncharted, there is growing consensus that CA2 pyramids do fire bursts of action potentials *in vivo* (Kay et al., 2016; Oliva, Fernandez-Ruiz, Buzsaki, & Berenyi, 2016) and increase their activity during social and aggressive encounters (Donegan et al., 2020; Leroy et al., 2018), but see also (Alexander et al., 2016)), consistent with the peptidergic modulation of CA2 firing described here.

## Supplemental Figures

**Figure 1 - Supplemental 1.**
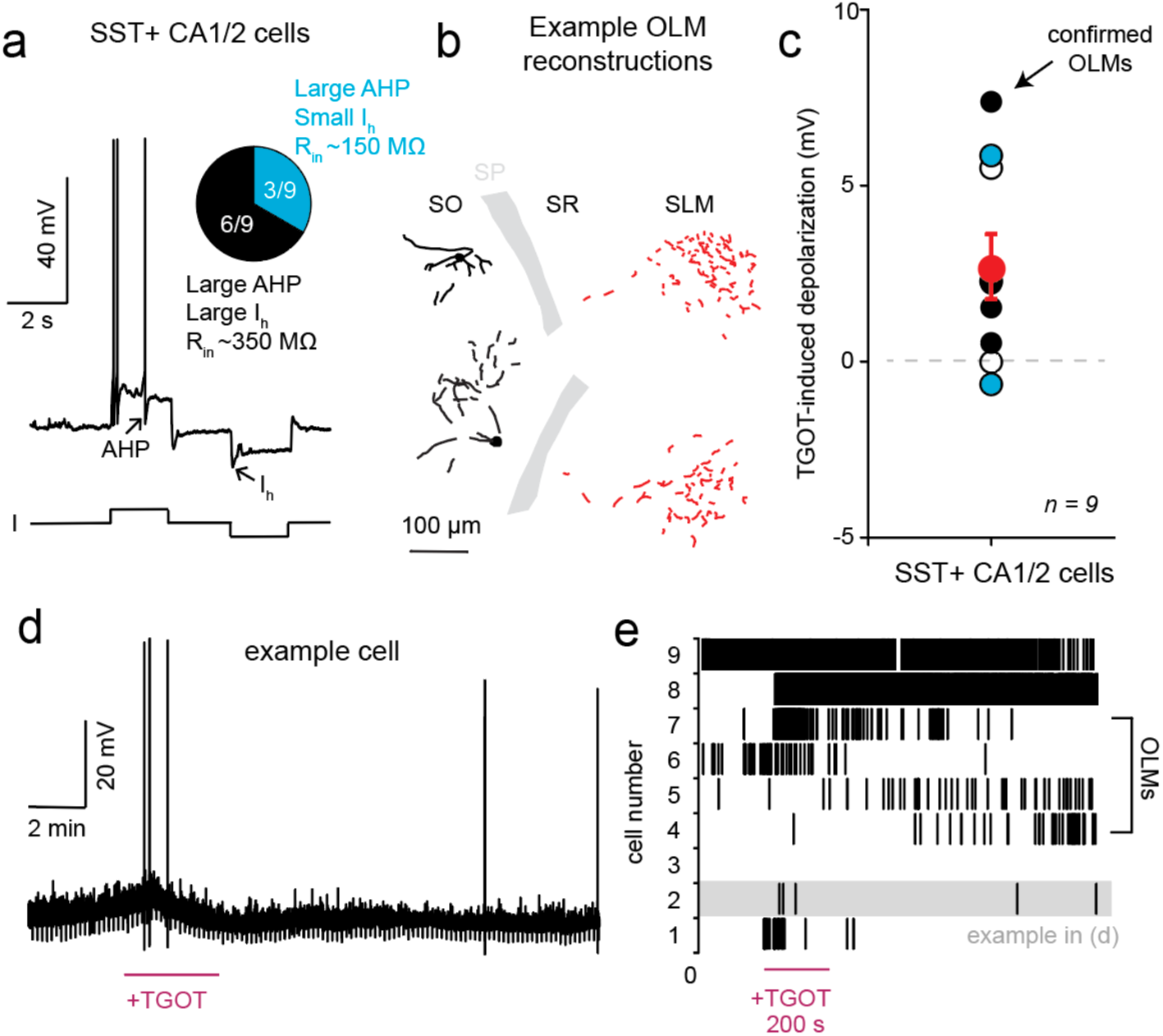
SST+ CA2 neurons show variable, small responses to OXTR stimulation. CA2 SST+ neurons were targeted using a transgenic reporter mouse line (SST-*ires*-Cre X Ai9) and characterized by electrophysiological properties like the after-hyperpolarization (AHP) and the sag current mediated by Ih (a) and morphological reconstructions (b). In (b) the soma and and dendrites are shown in black, while the axon is in red. Most SST+ neurons showed a mild depolarization in response to TGOT application (c). Group average is shown in red. (d) Example response in an SST+ neuron. Spiking responses were variable, as shown in a raster plot representing the group data in (e). Error bars reflect standard error of the mean. The 9 cells were recorded in 3 different mice. Related to Figure 1.

**Figure 2 - Supplemental 1.**
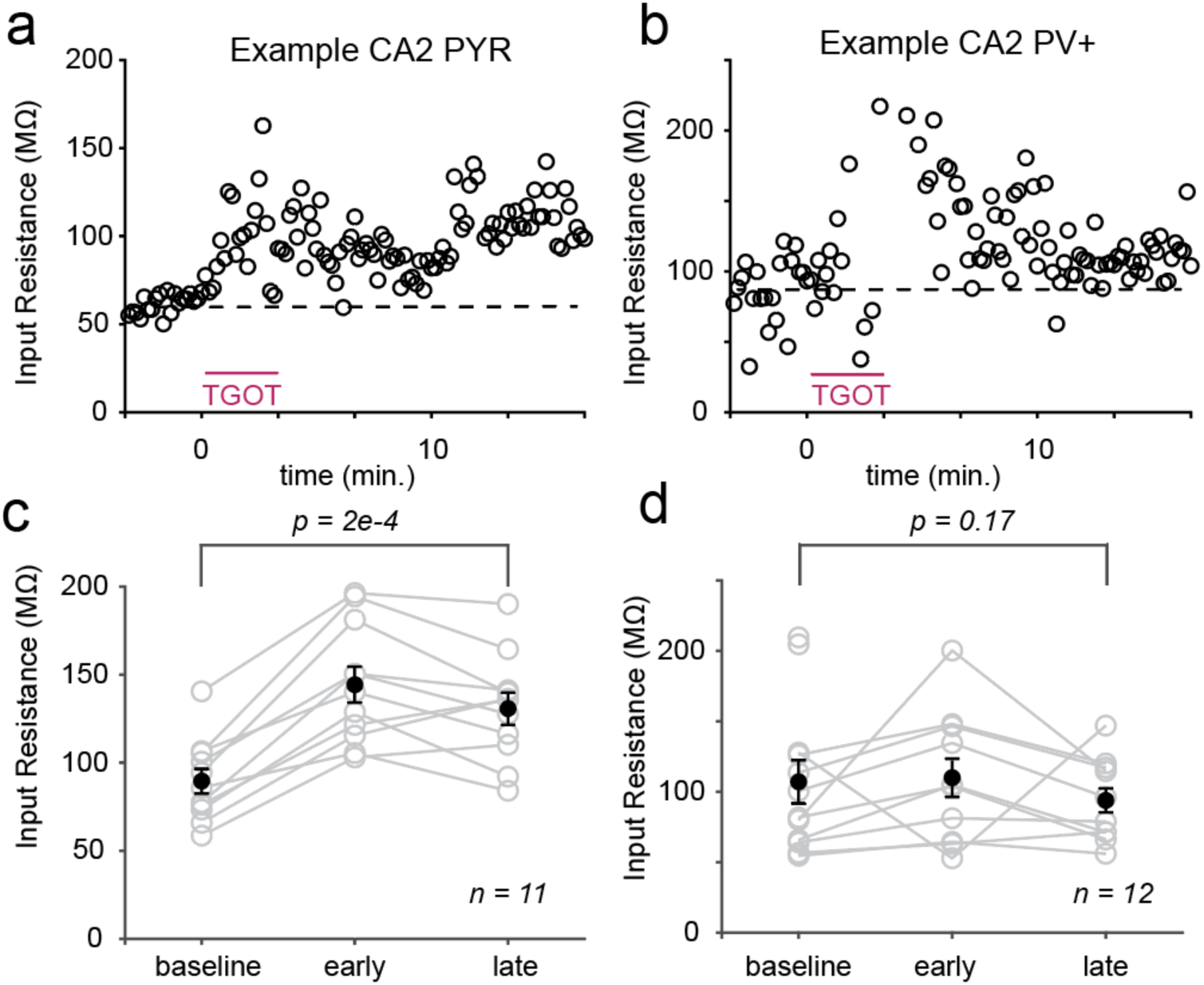
Upon OXTR stimulation, cellular input resistance increases, with different time courses, in CA2 PYRs and PV+ cells. Input resistance, measured every 10 seconds in an example CA2 pyramidal cell (a) and PV+ interneuron (b). A dashed line at baseline is added for visual clarity. Input resistance remains elevated, 15 minutes after drug application in CA2 PYRs (c), while it returns to baseline in PV+ cells (d). Group average is shown in red. Error bars reflect standard error of the mean. P-values are the result of paired *t*-tests. Data are from the same cells depicted in Figure 1. Related to Figure 2.

**Figure 3 - Supplemental 1.**
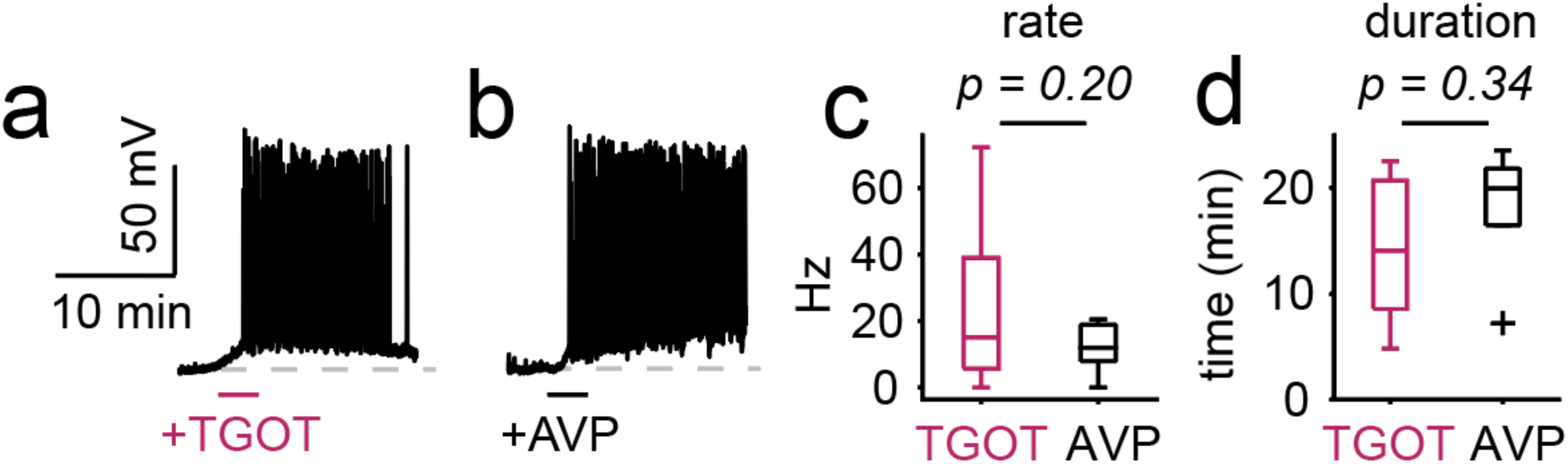
Oxytocin and vasopressin elicit comparable burst firing in CA2 pyramidal cells. Application of oxytocin (TGOT, 400 nM) and vasopressin (AVP, 1 μM) receptor agonists elicits depolarization and burst firing in CA2 PYRs (a, b). Average burst firing rate (c) and duration (d) are comparable between AVP (n = 7 cells / 4 mice) and TGOT (n = 18 cells / 12 mice). Related to Figure 3.

**Figure 3 - Supplemental 2.**
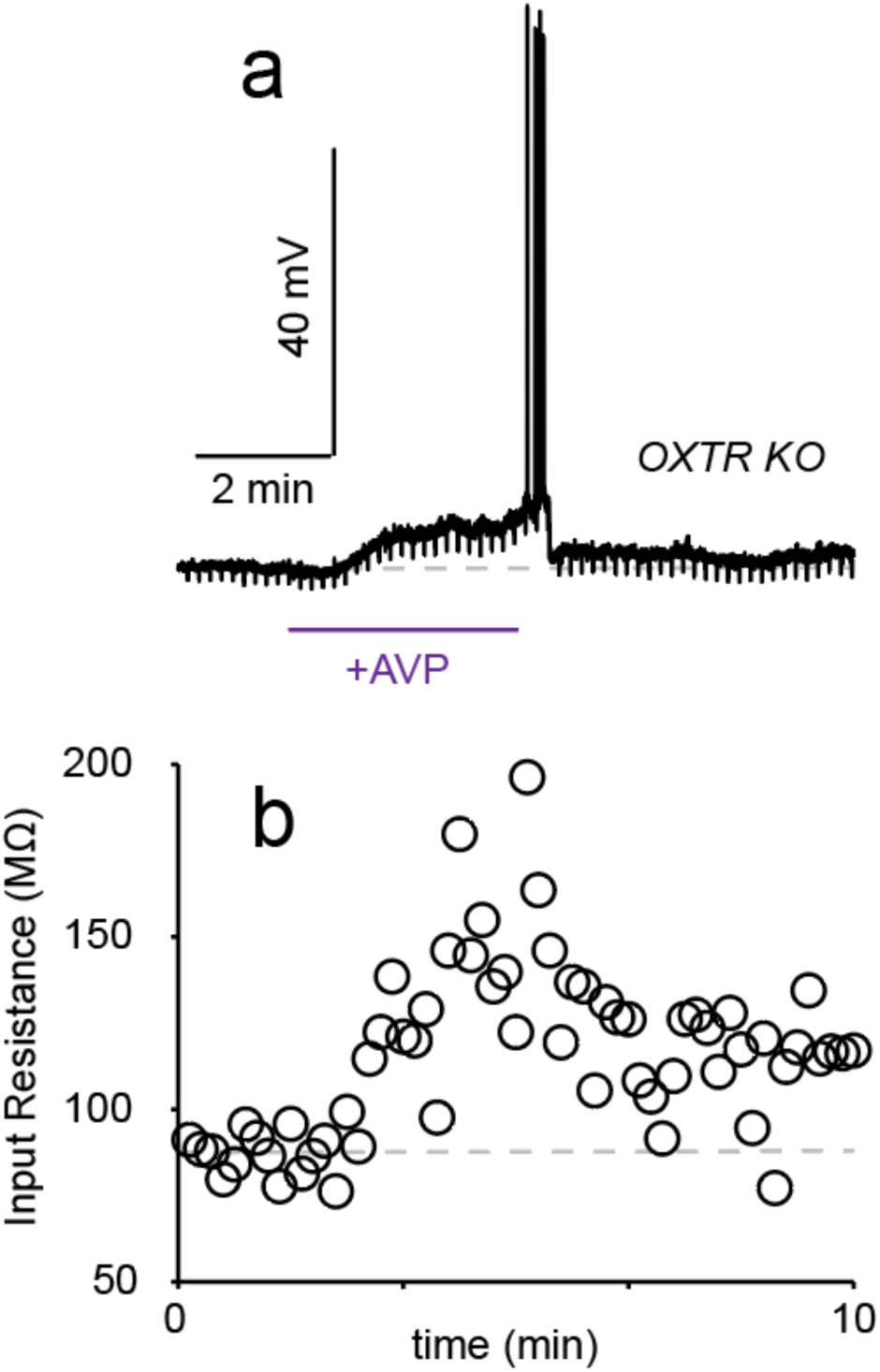
Vasopressin can excite CA2 pyramidal cells independent of the OXTR. Example recording from a CA2 PYR upon vasopressin (AVP) application in an OXTR KO animal (a). Accompanying input resistance measurements (b). Related to Figure 3.

**Figure 3 - Supplemental 3.**
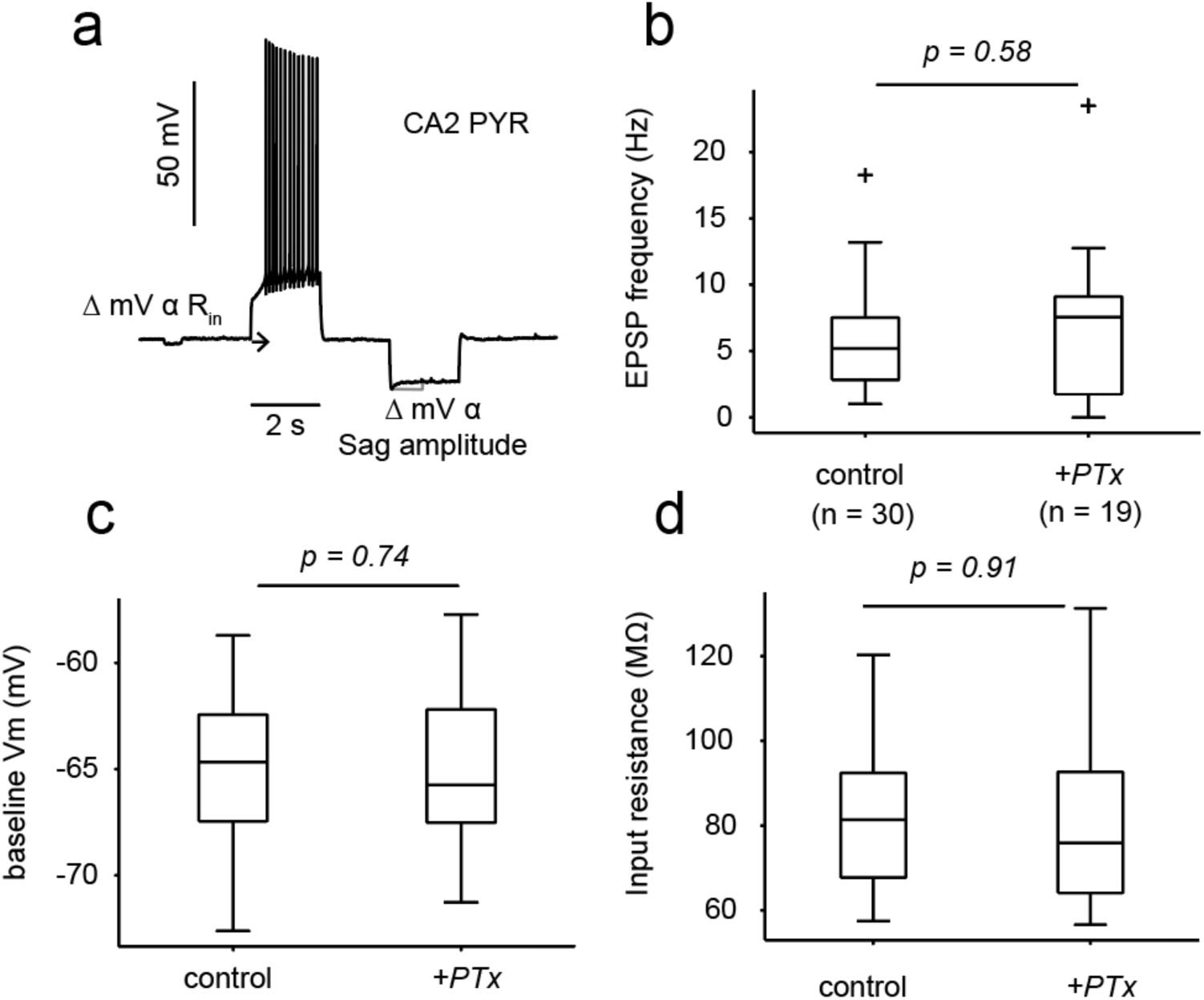
Pertussis toxin (PTx) pre-treatment (24-72 hours) does not alter intrinsic properties, of synaptic input, of CA2 PYRs. (a) Example voltage response in a CA2 PYR to the step protocol used to determine input resistance (R_in_) and sag ratio. (b) EPSP frequency recorded in CA2 PYRs pre-treated with PTx or in control slices. (c) Baseline membrane potential in CA2 PYRs with or without PTx pre-treatment. (d) Cellular input resistance, measured using a 20 pA, 500 ms hyperpolarizing step, in CA2 PYRs with or without PTx pre-treatment. P values are the result of unpaired *t*- tests. Related to figure 3.

**Figure 3 - Supplemental 4.**
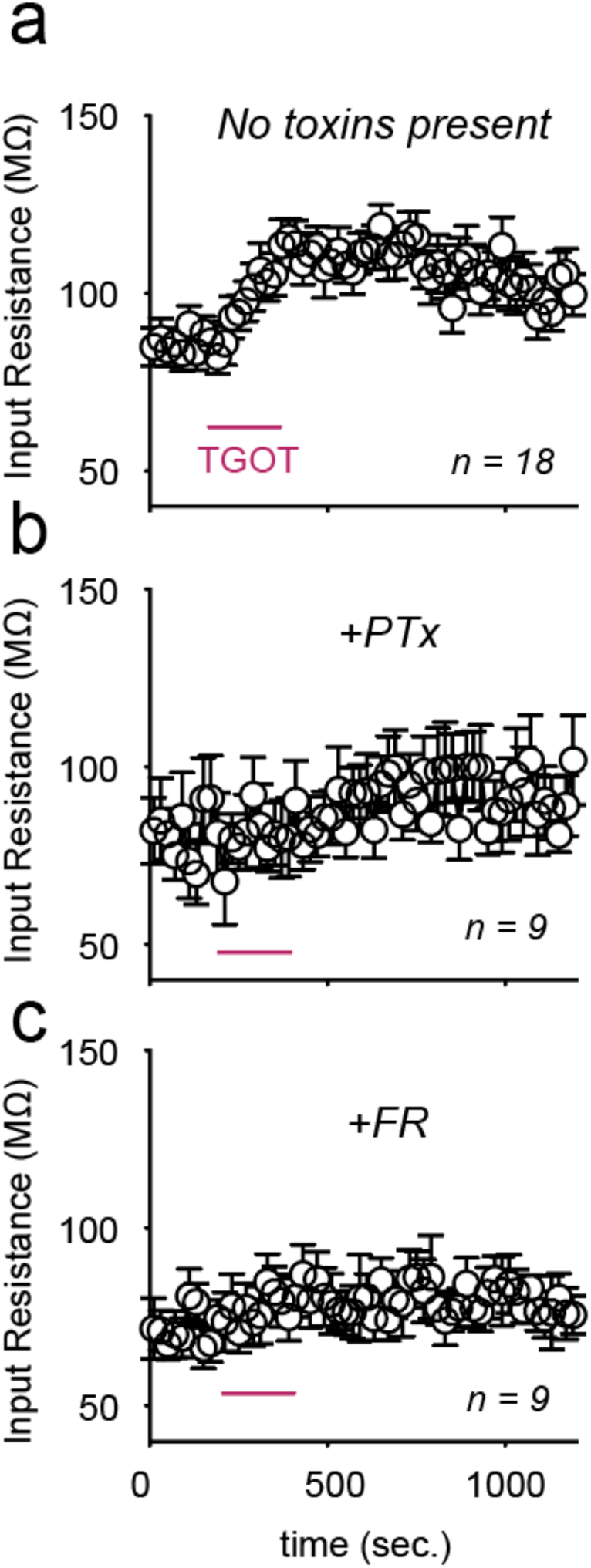
Pre-treatment with pertussis toxin (PTx) or FR900359 (FR) prevents TGOT-induced increase in input resistance. Input resistance measurements from CA2 PYRs upon TGOT application in control conditions (a), or given pertussis toxin (PTx) pre-treatment (b), or FR900359 (FR) pre-treatment (c). Related to figure 3.

**Figure 4 - Supplemental 1.**
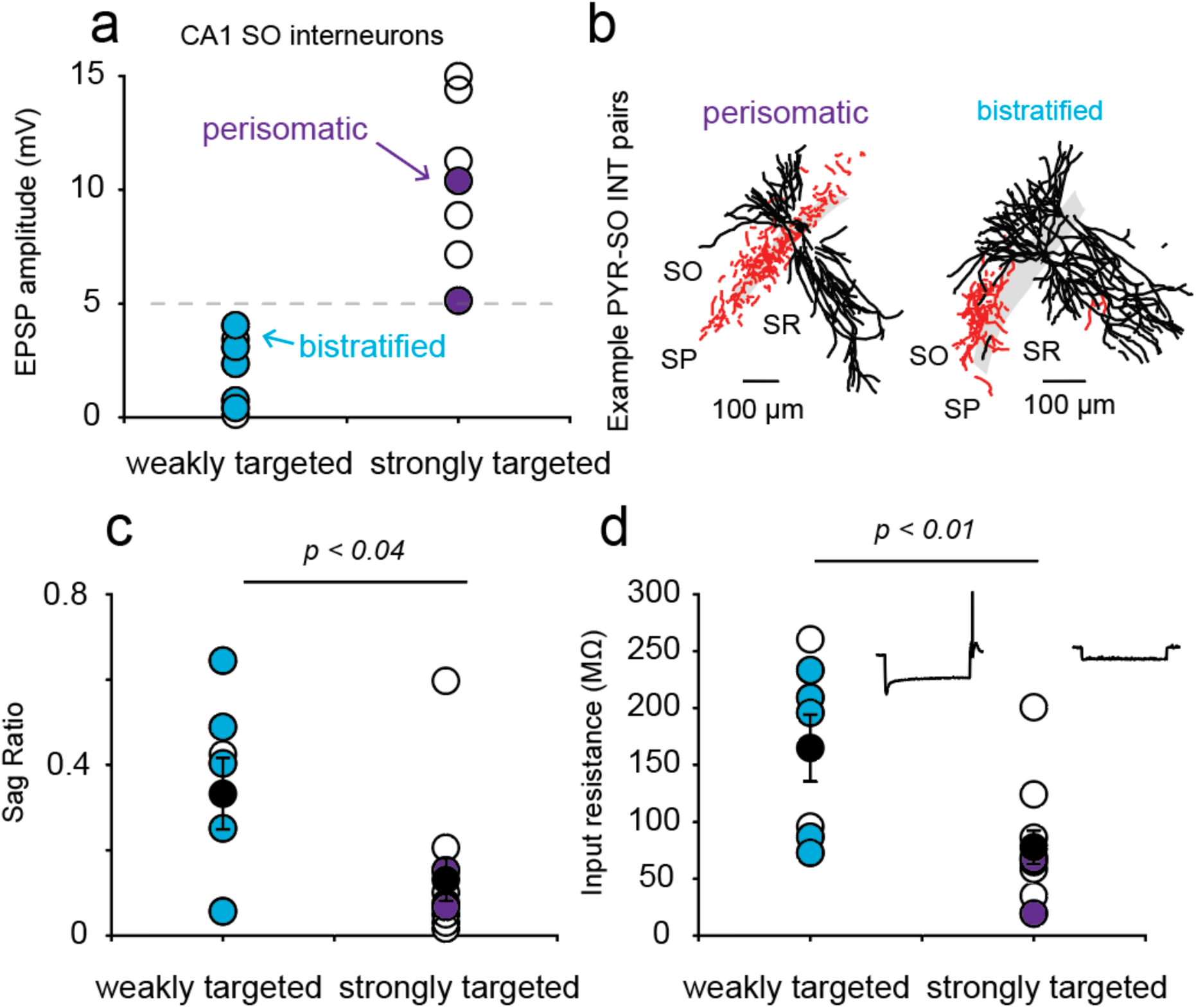
CA2 PYRs differentially target different interneuron subtypes. CA1 *stratum oriens* interneurons were grouped based on the amplitude of the EPSP evoked by CA2 PYR stimulation. Strongly targeted interneurons were defined as those with EPSPs of greater than 5 mV, while weakly targeted interneurons had EPSPs <5 mV (a). When possible, posthoc reconstructions were used to anatomically classify the interneurons recorded (b). Strongly targeted interneurons differed from those weakly targeted in the amount of “sag” produced by a hyperpolarizing pulse (relative to the steady state response; c) and input resistance (d). Example responses to hyperpolarizing current injection are shown in the insets. Cells with perisomatic (purple) or bistratified (blue) axonal arborizations are color coded in a-d. Related to figure 4.

**Figure 5 - Supplemental 1.**
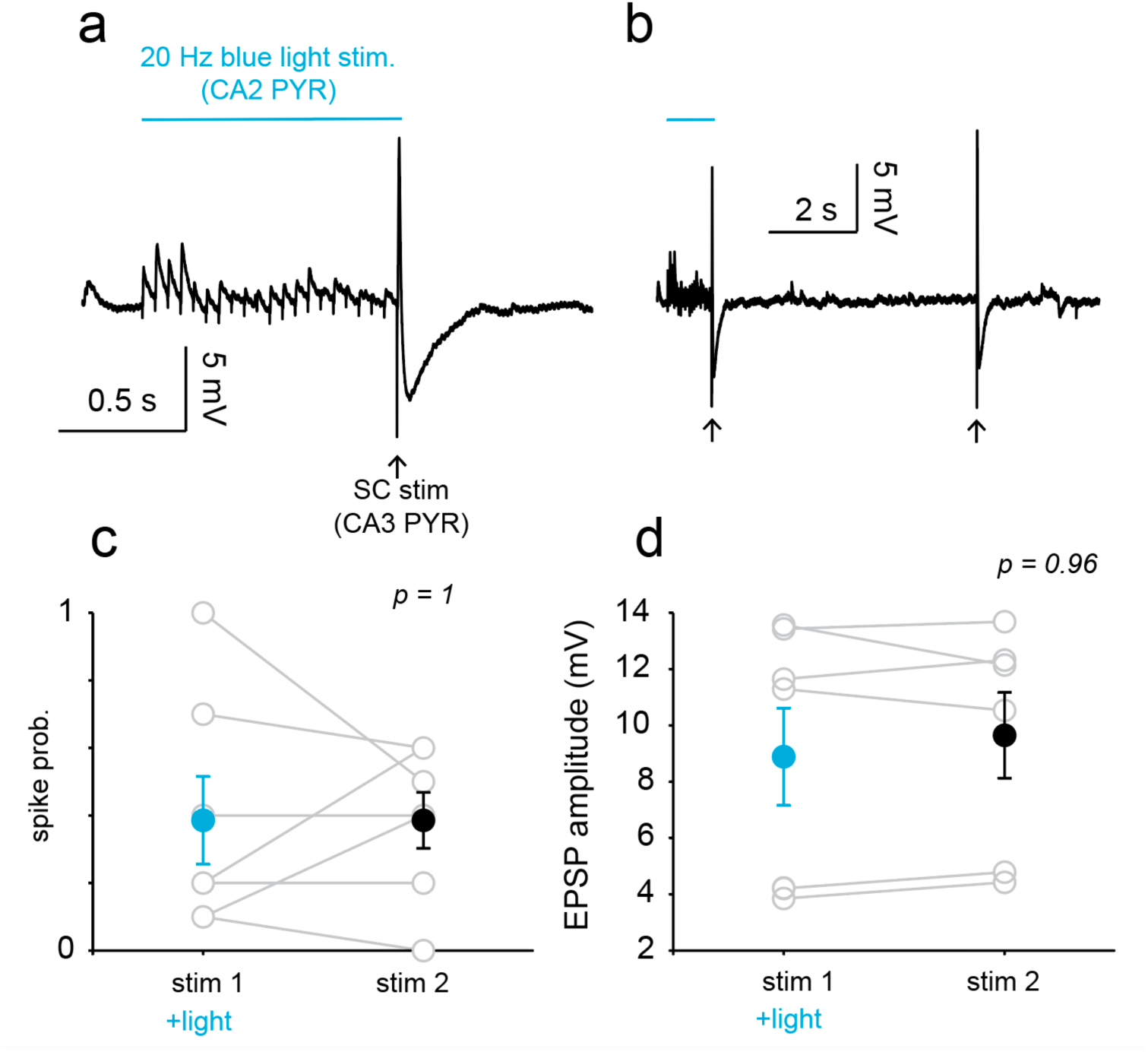
CA2 PYR bursts alone do not alter CA3- CA1 transmission. Current clamp recording from a CA1 PYR during optogenetic stimulation of CA2 PYRs (20Hz, 1 sec blue light in Amigo2-Cre mice injected with a floxed channelrhopsin variant) and electrical stimulation of the Schaffer collaterals (SC; a, b). Quantification of SC evoked spike probability (c) and EPSP amplitude (d) in the absence or presence of light stimulation. Both groups are statistically indistinguishable when analyzed using paired *t*-tests. Related to figure 5.

**Figure 5 - Supplemental 2.**
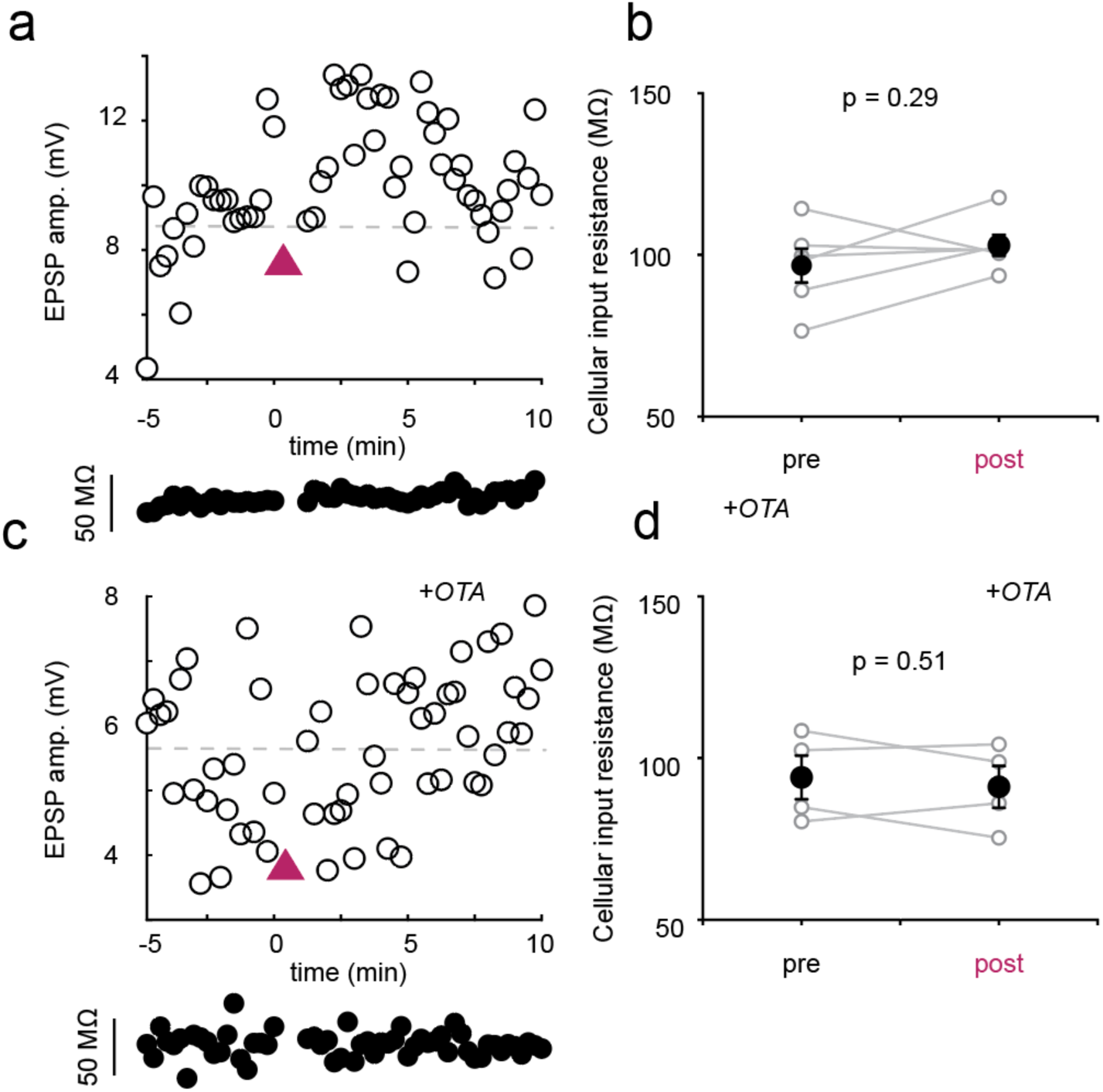
Cellular input resistance is stable during optogenetic recordings. (a) Schaffer collateral (SC) – evoked EPSP amplitude in an example CA1 PYR throughout the recording session. Input resistance, which is sampled throughout the recording, is shown below. The magenta triangle indicates the time of blue light stimulation. (b) Quantification of input resistance before and after optogenetic stimulation. Measurements of EPSP amplitude and input resistance in an example CA1 PYR pre-treated with the OXTR antagonist OTA (1 μM). (d) Group data from OTA-treated cells. P-values are the result of paired *t*- tests. Related to figure 5.

**Figure 5 - Supplemental 3.**
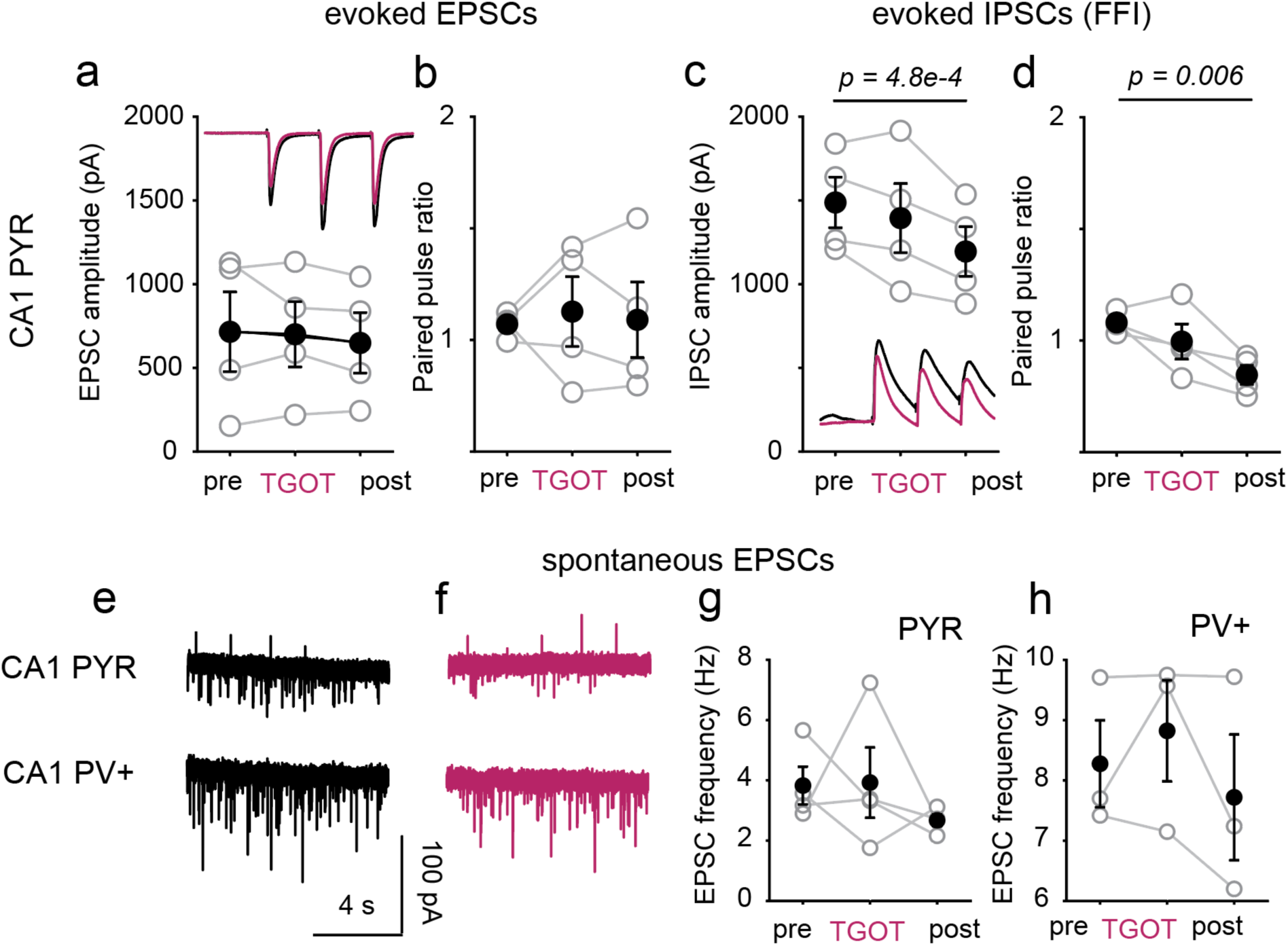
TGOT does not affect directly affect CA3-CA1 transmission. (a) Schaffer collateral evoked EPSC amplitude and paired-pulse ratio (b), before, during and 5 minutes after TGOT treatment. Inset shows data from an example cell. (c) Evoked feed-forward IPSCs and paired-pulse ratio (d) recorded in CA1 PYRs upon SC stimulation. (e) Spontaneous excitatory currents recorded in CA1 pyramidal cells (top) and PV+ interneurons (bottom) under control conditions and with TGOT present (f). Quantification of EPSC frequency across treatment conditions (g, h). P-values are the result of a paired *t*-test comparing pre and post conditions. PYR data from 4 cells / 2 mice. PV data from 3 cells / 2 mice. Related to Figure 5.

**Figure 5 - Supplemental 4.**
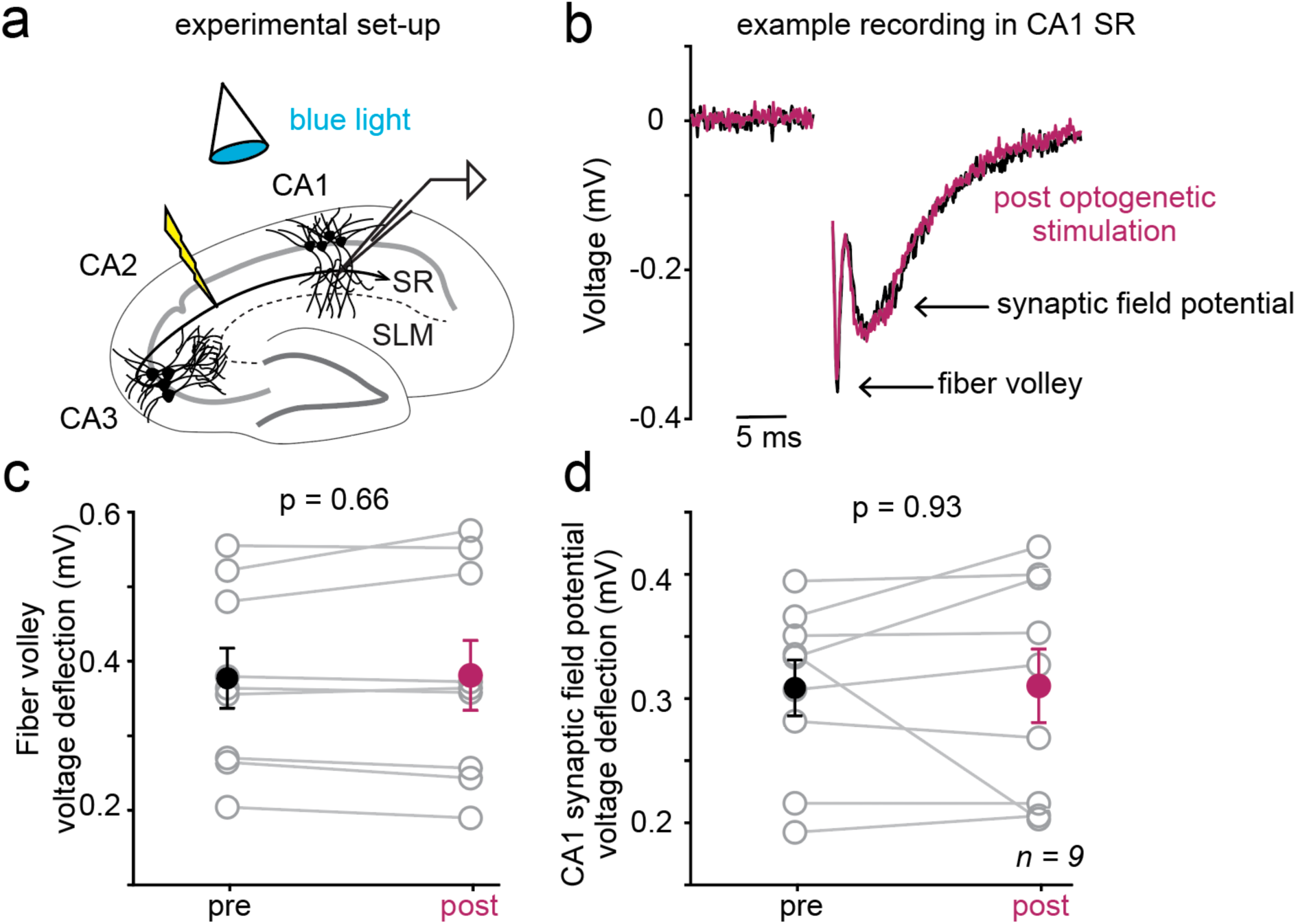
CA3 pyramidal cell fiber volley is unchanged upon endogenous oxytocin release. (a) Schematic depicting experimental set-up. CA3 axons are electrically stimulated near CA1 to avoid exciting CA2, while the Schaffer Collateral (SC) fiber volley and synaptic field potential were recorded in CA1 *stratum radiatum*. (b) Trial-averaged example recording from an exemplary cell before and after (magenta) optogenetic stimulation of oxytocinergic fibers. (c) Quantification of fiber volley deflection across cells and conditions. (d) Quantification of the CA1 synaptic field potential across cells and conditions. P values are the result of paired t-tests. We note that the SC-evoked synaptic field potential was unchanged upon optogenetic stimulation of oxytocinergic fibers, unlike the intracellularly recorded EPSP, likely due to differences in the stimulation configuration. In these experiments, the stimulation intensity was designed to elicit visually distinguished axonal and synaptic potentials and was smaller than used in other experimental configurations. Data from 9 cells / 4 mice. Related to figure 5.

## Methods

### Experimental model

All procedures involving animals were approved by the Institutional Animal Care and Use Committee at the New York University Langone Medical Center (NYULMC), and in accordance with guidelines from the National Institutes of Health. Animals were housed in fully equipped facilities in the Science Building, which is operated by NYULMC’s Division of Comparative Medicine. Male and female mice, post-natal days 50 - 90, were used in all experiments. No physiological differences were observed between sexes and data was pooled. Non-transgenic littermates were used as controls in experiments involving transgenic mouse lines. Homozygous *Oxytocin*-*ires*-Cre (Jackson Labs; Stock No. 024234), hemizygote *Amigo2-*Cre mice (Jackson Labs; Stock No. 030215) and homozygous *OXTR-ires-Cre* (provided by Dr. Katsuhiko Nishimori (Tohoku University)) were used for optogenetic studies. Targeted interneuron recordings were made from the offspring of crosses between either PV-Cre (Jackson Labs; Stock No. 008069) or SST- ires-Cre mice (Jackson Labs; Stock No. 013044) with Ai9 mice (Jackson Labs; Stock No. 007909).

### Stereotaxic injections

For all stereotaxic surgeries, mice (aged 4 – 10 weeks) were anesthetized with isofluorane (2%–5%) and secured in a stereotaxic apparatus (Kopf). Glass pipettes (Drummond Scientific) were formed using a P-2000 puller (Sutter Instrument) and were characterized by a long taper and 10-20 μm diameter tips. Pipettes were back-filled with mineral oil (Fisher Scientific) before being loaded with virus or toxin (Nanoject II, Drummond Scientific) and positioned at the stereotaxic coordinates indicated below. A small drill hole was made in the skull to allow for pipette insertion. To optogenetically excite cells, we injected pAAV5-EF1a-DIO-ChETA-eYFP-WPRE-HGHpa (Addgene). Details on each surgery are provided below:

| *Region* | *Lateral coordinates (mm)* | *Anterior (+) / Posterior (-) coordinates (mm)* | *Dorsal (+) / Ventral (-) coordinates (mm)* | *Volume and rate* | *Notes* |
| --- | --- | --- | --- | --- | --- |
| Paraventricular nucleus (PVN) of the hypothalamus | 0.25 | -0.7 | -4 | 69 nL of virus every 30 s for 5 minutes (690 nL / hemisphere) | Bilateral injections into Oxytocin-ires-Cre mice; 3-4 weeks incubation |
| CA2 subregion of the hippocampus | 1.6 | -1.6 | -1.7 | 23 nL of virus every 20 s for 80 s (115 nL / hemisphere) | Bilateral injections into Amigo2-Cre and OXTR-ires-Cre mice ; 2 weeks incubation |
| Lateral ventricle | 1 | 0.3 | -3 | 1 – 2 μL | Unilateral injection of pertussis toxin (0.1 g/L; 24 – 72 hours incubation) |

Throughout the surgery, body temperature, breathing and heart rate were monitored. Saline was administered subcutaneously (s.c) to maintain hydration and the animal was monitored post-operationally for signs of distress and discomfort. Buprenorphine (0.1 mg/kg, s.c) was given for analgesia. Successful targeting of viral constructs was confirmed via posthoc epifluorescence imaging (Zeiss LSM 510 Imager.M1 confocal microscope). Photostimulation of ChETA was achieved through 470 nm light delivered to the slice field through a 5x or 40x objective. Illumination intensity was adjusted between 0.1-2.0 mW, depending on the experimental intention.

### Electrophysiology and recordings

Following anesthesia induced with a mixture of ketamine/xylazine (150 mg/kg and 10 mg/kg, respectively), adult mice were transcardially perfused with oxygenated, ice-cold sucrose solution containing (in mM): 206 Sucrose, 11 D-Glucose, 2.5 KCl, 1 NaH_2_PO_4_, 10 MgCl_2_, 2 CaCl_2_ and 26 NaHCO_3_. Following perfusion and decapitation, brains were removed and placed in the cold sucrose solution for sectioning (Leica VT 1000S Vibratome). Transverse, 300 - 350 μm bilateral hippocampal sections were cut and transferred to an oxygenated, 34° C recovery chamber filled with artificial cerebro-spinal fluid (aCSF) containing (in mM): 122 NaCl, 3 KCl, 10 D-Glucose, 1.25 NaH_2_PO_4_, 2 CaCl_2_, 1.3 MgCl_2_, and 26 NaHCO_3_. Slices recovered for 30 minutes at 34° C before they were transferred to room temperature for 1 - 4 h before recording.

Slice recordings were performed in a submerged chamber maintained at approximately 34° C with constant bath perfusion of aCSF at ∼2 mL/min. Whole cell recordings were made with borosilicate glass pipettes (2 – 4 MΩ) pulled on a Sutter Instrument P-97 micropipette puller. The same intracellular solution was used for current and voltage clamp experiments and contained (in mM): 130 K-Gluconate, 1 MgCl_2_, 10 HEPES, 0.3 EGTA, 10 Tris-Phosphocreatine, 4 Mg-ATP, and 0.3 Na-GTP. Biocytin (0.1%) was including in the internal solution for morphological reconstruction of recorded cells. Hippocampal regions and layers were identified visually with an upright microscope (Zeiss Axioskop 2 FS Plus) using infrared differential interference contrast optics. CA2 pyramidal cells were identified by their distinct electrophysiological properties (Chevaleyre & Siegelbaum, 2010; Tirko et al., 2018). Recordings were made using a MultiClamp 700B amplifier (Axon Instruments, Union City, CA). Signals were filtered at 10 kHz using a Bessel filter, digitized at 20kHz with a Digidata 1322A analog-digital interface (Axon Instruments) and analyzed using custom MATLAB scripts (MathWorks). Cellular input resistance was monitored, every 10 seconds, throughout most recordings by regularly giving a small hyperpolarizing step. Negative input resistance values, and those that were more than 2.5 times away from the baseline value were omitted.

Each cell represented an independent biological replicate. Recordings were excluded from further analysis if significant swings in series resistance (>25% change in voltage clamp experiments or a visible shrinkage of the action potential amplitude in current clamp recordings) or membrane potential (during the first few minutes of recording) were observed). Similarly, recordings were not included in analysis if a stable recording would not be obtained after drug or electrical intervention. Whenever possible, littermates were used in experimental and control groups. Collection of control data was interleaved with collection of experimental data to minimize any effects of experimental drift over time.

For experiments involving channelrhodopsins, viral expression was confirmed by expression of YFP in the hippocampus or hypothalamus before recordings began.

Electrical stimulation of the Schaffer Collaterals was achieved by placing a Tungsten microelectrode (A&M Systems) in the *stratum radiatum* layer of CA2.

### Fiber Volley

Patch pipettes with a resistance of 4∼6 MΩ were made from borosilicate glass (World Precision Instruments) with a Sutter Instrument P-97 micropipette puller and filled with 1M NaCl solution. The stimulation electrode was made from the same borosilicate glass pulled into a tapered tip, trimmed with scissors, and filled with aCSF. Both recording and stimulation electrodes were placed in the SR of CA1 with the stimulation electrode closer to the CA2/CA1 border. Electrical stimulation was performed using a model 2100 Isolated Pulse Stimulator (A-M Systems). All data were sampled and analyzed using Clampfit 10.2 software (Molecular Devices) and MATLAB (MathWorks). Evoked synaptic potentials were recorded under passive current clamp with Gain = 50. A unipolar stimulus with duration 0.01 to 0.1ms was used to elicit the fiber volley signal. Field potentials were recorded during incremental increase of stimulus strength from 0 until the fiber volley signal size saturated or merged into stimulation artifact to generate the input/output curve for each slice. Then the stimulus strength was returned to the intensity that produced, approximately, the half-maximal response and maintained with an inter-stimulus interval of 20s. After at least 10 min of baseline control recording, a prolonged blue light stimulation (20Hz, 80s) was applied to induce endogenous neuropeptide release from PVN axonal terminals expressing ChETA.

### Immunohistochemistry

In preparation for imaging and biocytin reconstruction, slices were transferred from the recording chamber to 4% paraformaldehyde in phosphate buffered solution (PBS, Affymetrix) overnight. After washing with PBS with 0.1% Tween 20 (PBST, Sigma), the slices were left in 30% sucrose in PBST for at least 48 hours. Streptavidin-647 (1:350 dilution, Molecular Probes) was used to visualize recorded cells and was applied for 2 hours at room temperature before washing and mounting. Images were acquired on a Zeiss LSM 510 Imager.M1 confocal microscope and tracing for 2D morphological reconstruction was performed using NeuroLeucida software.

Key resources used in this paper:

| REAGENT or RESOURCE | SOURCE | IDENTIFIER |
| --- | --- | --- |
| TGOT ((Thr<sup>4</sup>,Gly<sup>7</sup>)-Oxytocin; 400 nM) | Bachem | Cat# 4013837 |
| Retigabine (100 µM) | Alomone | Cat# R-100 |
| Pertussis toxin | Sigma | Cat# P7208 |
| FR900359 / UBO-QIC (1 µM) | Drs. Evi Kostenis and Gabriele König (University of Bonn) | N/A |
| **Virus Strains** |  |  |
| pAAV5-EF1α-DIO-ChETA-eYFP | Penn Vector Core | N/A |
| **Experimental Models: Organisms/Strains** |  |  |
| Mouse: C57B/6J | Jackson Labs | Cat# 000664 |
| Mouse: B6;129S-*Oxt*<sup>tm1.1(cre)Dolsn</sup>/J (*Oxt-ires-cre*) | Jackson Labs | Cat# 024234 |
| Mouse: B6.Cg-Tg(*Amigo2-cre*)1Sieg/J (*Amigo2-Cre*) | Jackson Labs | Cat# 030215 |
| Mouse: *Oxtr* cDNA<sup>HA</sup>-*Ires-Cre* (*OXTR-Cre*) | Drs. Katsuhiko Nishimori and Shizu Hidema (Tohoku University) | N/A |
| Mouse: B6;129P2-*Pvalb*<sup>tm1(cre)Arbr</sup>/J (*PV-Cre*) | Jackson Labs | Cat# 008069 |
| Mouse: *Sst*<sup>tm2.1(cre)Zjh</sup>/J (*Sst-Cre*) | Jackson Labs | Cat# 013044 |
| Mouse: B6.Cg-Gt(ROSA)26Sor<sup>tm9(CAG-tdTomato)Hze/J</sup> (Ai9) | Jackson Labs | Cat# 007909 |

Data generated in this work are available on Dryad:

Eyring, Katherine et al. (2020), Oxytocin signals via Gi and Gq to drive persistent CA2 pyramidal cell firing and strengthen CA3-CA1 neurotransmission, Dryad,

Dataset, https://doi.org/10.5061/dryad.0vt4b8gw6

## Acknowledgements

We are grateful to Dr. Steven Siegelbaum for generously providing the Amigo2-cre mice. We thank Vincent Robert for expert technical advice and Guoling Tian for excellent technical assistance.We thank Tsien lab members and Drs. Jayeeta Basu, Brian Kobilka, Bertil Hille and Mark Shapiro for helpful discussions.

## Competing Interests

The authors declare no competing interests.

